# Dissecting lipid contents in the distinct regions of native retinal rod disk membranes

**DOI:** 10.1101/2021.01.11.426250

**Authors:** Christopher L. Sander, Avery E. Sears, Antonio F. M. Pinto, Elliot H. Choi, Shirin Kahremany, Hui Jin, Els Pardon, Susie Suh, Zhiqian Dong, Jan Steyaert, Alan Saghatelian, Dorota Skowronska-Krawczyk, Philip D. Kiser, Krzysztof Palczewski

## Abstract

Photoreceptors rely on distinct membrane compartments to support their specialized function. Unlike protein localization, identification of critical differences in membrane content has not yet been expanded to lipids, due to the difficulty of isolating domain-specific samples. We have overcome this by using SMA to co-immunopurify membrane proteins and their native lipids from two regions of photoreceptor ROS disks. Each sample’s copurified lipids were subjected to untargeted lipidomic and fatty acid analysis. Extensive differences between center (rhodopsin) and rim (ABCA4 and PRPH2/ROM1) samples included a lower PC to PE ratio and increased LC- and VLC-PUFAs in the center relative to the rim region, which were enriched in shorter, saturated FAs. The comparatively few differences between the two rim samples likely reflect specific protein-lipid interactions. High-resolution profiling of the ROS disk lipid composition provides a model for future studies of other complex cellular structures, and gives new insights into how intricate membrane structure and protein activity are balanced within the ROS.

**SUMMARY:** Sander et al. have parsed the lipid composition of native-source photoreceptor disks and find large differences in fatty acid unsaturation and chain length between the center and rim regions. They selectively copurify membrane proteins and lipids from each region in SMALPs using nanobodies and antibodies.

## INTRODUCTION

Photoreceptor cells of the retina are highly differentiated neurons responsible for the capture of light and conversion of its energy to the biochemical amplification cascade known as phototransduction. Each of these cells has a highly elongated cilium called the rod outer segment (ROS), which is composed of an internal stack of membranous disks surrounded by plasma membrane. Roughly 40 million molecules of rhodopsin are packed into each ROS, and each light-activated rhodopsin is capable of binding and activating many molecules of transducin (the G protein of the visual system) (Fung et al., 1981; Nathans, 1992; Polans et al., 1996; Heck and Hofmann, 2001). To accommodate the uniquely dynamic, yet exquisitely structured environment, ROS disks contain specialized lipids rich in long chain and very long chain polyunsaturated fatty acids (LC-PUFAs and VLC-PUFAs, respectively). The disks are also known to have significantly higher levels of phosphatidylethanolamine (PE) than are typically found in plasma membranes. This overabundance of PE is compensated by a relative scarcity of phosphatidylcholine and phosphatidylserine (PC and PS, respectively) in ROS membranes (Boesze-Battaglia and Albert, 1992). Cholesterol has also been found to be necessary for rhodopsin activity; however, excessively high concentrations of cholesterol reduce its signaling (Mitchell et al., 1990, 1992a). Indeed, many components of the membrane can have a profound impact on the function of the membrane proteins therein, making high-resolution study of membrane environments critical to the overall characterization of membrane proteins. Early work by Falk and Fatt on the ultra-structure of ROS membranes showed a remarkable ability of the outer rim region of ROS disks to resist disruption in the presence of Tris buffer after OsO_4_ fixation (Falk and Fatt, 1969). Their work gives an indication that the membranes in the rim region are distinct from the center, but concrete evidence in support of this has not yet been offered.

The current lack of knowledge regarding molecular differences between the center and rim of ROS disk membranes represents a significant bottleneck in the study of lipid synthesis, metabolism, and transport (Zhang et al., 2001; Edwards et al., 2001; Chen et al., 2005, 2007; Berdeaux et al., 2010; Sapieha et al., 2011; Chen et al., 2013, 2020). These processes modulate the impact of lipids on retinal degenerative diseases, such as Stargardt-like macular dystrophy type 3, retinitis pigmentosa, diabetic retinopathy, and age-related macular degeneration (Simonelli et al., 1996; Bernstein et al., 2001; Seddon, 2003, 2006; SanGiovanni et al., 2007; Liu et al., 2010; Tikhonenko et al., 2010, 2013; Logan et al., 2013; Logan and Anderson, 2014; Hiebler et al., 2014). Mapping the possible lipid domains in which vision-related membrane proteins reside would be an invaluable contribution to the study of protein-lipid interactions.

The coupled processes of phototransduction and the visual cycle utilize several membrane proteins. Rhodopsin, the prototypical G protein-coupled receptor (GPCR) responsible for initiating phototransduction, appears to prefer limited cholesterol content for optimal activity (Mitchell et al., 1992b; Palczewski, 2006). Rhodopsin also exhibits maximal activity in a phospholipid environment with a high proportion of docosahexaenoic acid (DHA, 22:6) (Mitchell et al., 1992b). In the rim of ROS disks, ATP-binding cassette protein, family A, number 4 (ABCA4) assists in chromophore clearance from the ROS disk through *N*-retinylidene-phosphatidylethanolamine flippase activity. Prior work has shown that this lipid and all-*trans* retinal flippase is optimally active when the membrane contains at least 40% PE; it shows no activity in pure phosphatidylcholine (PC) liposomes (Sun and Nathans, 2001; Quazi and Molday, 2013; Quazi et al., 2012). Such high levels of PE are common in the native disk membranes of ROS (Daemen, 1973). The peripherin2-ROS membrane protein 1 (PRPH2/ROM1) complex is also found in the rim of ROS disks (Molday et al., 1987). PRPH2/ROM1 does not exhibit enzymatic activity, but is an essential component of disk morphogenesis and maintenance of the curved, bulbous rim of ROS disks (Goldberg and Molday, 1996a; b; Loewen and Molday, 2000; Kevany et al., 2013; Zulliger et al., 2018; Milstein et al., 2020).

The recent advent of styrene-maleic acid lipid particles (SMALPs) has made it possible to extract the membrane bilayer into discrete disks containing the proteins therein (Knowles et al., 2009; Jamshad et al., 2011). However, there remains a question regarding the nativity of SMALP-encased membranes; *i.e*., do lipids “copurified” with native proteins represent the environment from which the protein was extracted? Accordingly, there have been recent reports on the lipid exchange dynamics of polymer-bound lipid nanodiscs (Cuevas Arenas et al., 2017; Schmidt and Sturgis, 2018; Danielczak and Keller, 2018). These studies showed that phospholipids extracted in SMALPs and diisobutylene maleic acid lipid particles (DIBMALPs) exchange rapidly, orders of magnitude faster than in large unilamillar vesicles (LUVs) or membrane scaffold protein (MSP) nanodiscs. These findings suggest that native membrane proteins, once extracted by SMA, might reside in a lipid environment that reflects the average lipid environment of the extracted tissue.

ROS disks provide an interesting test case for the potential rearrangement of lipids around native proteins extracted by SMA, because the disk rim contains a different protein population than the disk center. It is reasonable to expect that the center of the ROS disk maintains a unique environment, as rhodopsin is thought to pack in a paracrystalline manner while requiring ample free volume in the membrane for high levels of signaling (Mitchell et al., 1992b; Fotiadis et al., 2004). The ROS rim, however, needs to adopt and maintain a curved and bulbous structure. To investigate whether the lipid exchange rate in the case of mammalian native tissue is slow enough to enable retention of the local lipid environment, we chose to purify ABCA4 and rhodopsin from SMA-extracted ROS. Different copurifying lipids in each purified protein sample would suggest retention of the native membrane environments in the respective SMALPs. To confirm the membranes are native in composition and not the result of *post hoc* segregation, we purified an additional protein complex, PRPH2/ROM1, from the rim region, with the hypothesis that if SMALPs retain native membrane environments, we should observe similar lipid profiles between rim samples and conserved differences between rim and central samples.

Here, we report the isolation of the specific lipid environments surrounding rhodopsin, ABCA4, and the PRPH2/ROM1 complex (**Fig. 1, a and b**). The membrane composition analysis specific to these proteins will facilitate our understanding on the effect of membrane components on the protein function.

**Figure 1.**
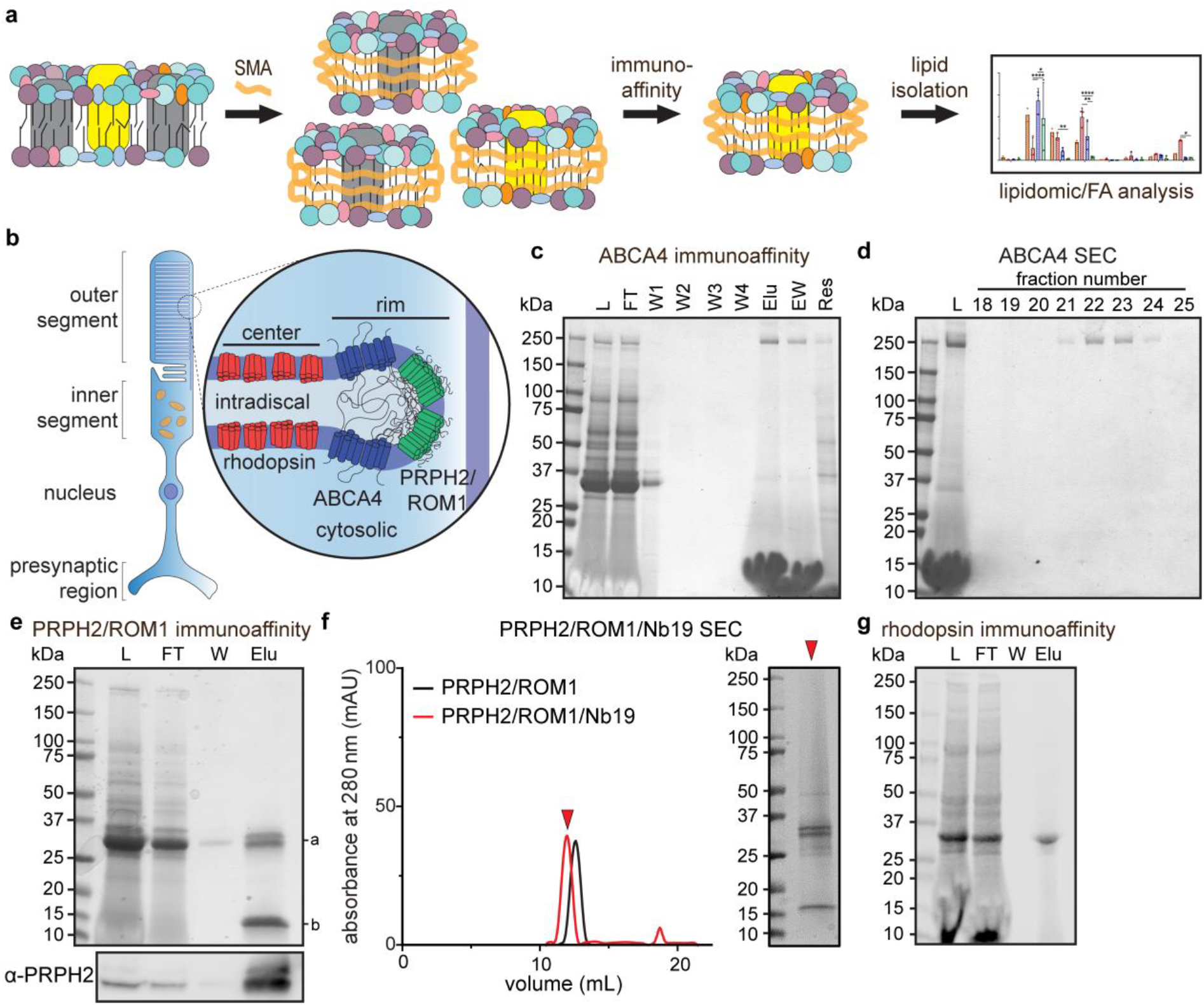
Detergent-free purification of native proteins from bovine ROS by immunoaffinity chromatography. (**a**) Native lipids isolated by the SMALP co-immunopurification procedure. SMA extracts membrane proteins with their native lipids; the SMALPs may then be subjected to immunoaffinity chromatography for purification of native nanodiscs, enabling analysis of copurifying lipids. (**b**) The intricate membrane structure of ROS disks in rod photoreceptors. Three major membrane protein components are rhodopsin, ABCA4, and PRPH2/ROM1. (**c**) Detergent-free, immunoaffinity purification of ABCA4 using the CL2 mAb. L, soluble ROS (16 mL, 10 µL loaded); FT, flow-through (16 mL, 10 µL loaded); W1-4, washes 1-4 (each 15 mL, 10 µL loaded); Elu, elution (1 mL, 10 µL loaded); EW, wash of column after elution (1 mL, 10 µL loaded); Res, resin (1 mL, 10 µL loaded). Stained with Coomassie Blue R250. (**d**) Detergent-free size exclusion chromatography (SEC) of combined elution fractions, 18-25 fraction numbers, 0.5 mL fractions from SEC, 10 µL loaded per lane. Stained with Coomassie Blue R250. (**e**) Detergent-free, immunoaffinity purification of PRPH2/ROM1 (a) using the Nb19 nanobody (b). L, soluble ROS (10 mL, 10 µL loaded); FT, flow-through (10 mL, 10 µL loaded); W, wash (10 mL, 10 µL loaded); Elu, elution (2.5 mL, 2.5 µL loaded). Bottom panel, anti-PRPH22 immunoblot of the above samples. (**f**) Detergent-free size exclusion chromatography of combined elution from Nb19-immunoaffinity purification. (*Left*) PRPH2/ROM1 incubated with Nb19 (red) elutes earlier than PRPH2/ROM1 alone (black). (*Right*) Peak PRPH2/ROM1/Nb19 fraction run on SDS-PAGE and stained with Coomassie Blue R250. (**g**) Detergent-free, immunoaffinity purification of rhodopsin using the 1D4 mAb. L, soluble ROS (16 mL, 10 µL loaded); FT, flow-through (16 mL, 10 µL loaded); W, wash (15 mL, 10 µL loaded); Elu, elution (1 mL, 10 µL loaded).

## RESULTS

### SMA extraction of ROS membrane proteins and development of mAb against ABCA4

We began by searching for a method of copurifying lipids in the immediate vicinity of each representative ROS membrane protein (**Fig. 1**). SMA showed a strong capacity for solubilizing ROS membrane proteins (**Fig. 1 a, Fig. S1 a**). The higher yield of total protein obtained from ROS extracted in SMA also showed near-complete extraction of the available ABCA4, as shown by immunoblot analysis (**Fig. S1 b**). Optimum extraction of ABCA4 in SMA occurred at 2.5% w/v and was essentially complete; in contrast, 2% LMNG (roughly 2000x the CMC) resulted in roughly 50% solubilization.

The C-terminal region of ABCA4 is an accessible site that contains a high affinity binding epitope for the Rim3F4 antibody (YDLPLHPRT) (Illing et al., 1997). The Rim3F4 antibody has very good affinity for the C-terminus of ABCA4, but immunopurification of ABCA4 proved difficult, given the low efficiency of elution from the column. We designed the CL2 monoclonal antibody to overcome low immunoaffinity efficiency by targeting an expanded region of the C-terminus. To develop a novel epitope near this site on the C-terminus, a 26 amino acid peptide (NETYDLPLHPRTAGASRQAKEVDKGC) from the near-extreme end of bovine ABCA4 (with the addition of a C-terminal cysteine) was synthesized and conjugated to keyhole limpet hemocyanin (KLH) protein to induce an immunogenic response in mice. **Fig. S1 c** shows the location and length of the resultant antibody binding site for CL2, in comparison to the locations of the antibody binding sites for Rim3F4 and TMR4 (Zhang et al., 2015), another antibody that targets the second extracytosolic domain.

Dot blot analysis of CL2 was conducted to determine whether the paratope was different from that of Rim3F4 (**Fig. S1 d**). Various cleavage products of the polypeptide, made by sequentially omitting two amino acids from each end, were adsorbed onto the membrane and then probed for CL2 binding. Compared to the full-length peptide, none of the putative sub-epitopes bound CL2 with nearly the same affinity. When the first residues were removed (ΔF2-6), there was a complete loss of binding, suggesting that they are integral to CL2 recognition. The affinities of those peptides missing the last few residues (ΔL2-6) were much weaker than that of the full-length sequence, indicating that both ends of the epitope are important for robust binding of CL2, and that CL2 uses a different epitope than Rim3F4. CL2 showed a relative lack of signal in the immunoblot of solubilized bovine ROS (**Fig. S1 f**). In comparison to Rim3F4 and TMR4, CL2 showed a weak but specific signal for ABCA4.

The relatively weak binding of CL2 was also apparent in murine samples (**Fig. S1 e and g**). The immunohistochemical analysis of murine retinas showed a gradual increase in ABCA4 signal intensity in samples stained with higher levels (lesser dilutions) of CL2 (**Fig. S1 e**). This staining profile contrasted with the profile for Rim3F4, which showed a robust signal in the outer segments even with the greatest dilution. CL2 showed a level of signal comparable to that of Rim3F4 for the same murine samples *via* immunoblots; and also showed comparable specificity, with no ABCA4 detected by CL2 or Rim3F4 in isolated, murine RPE, although small amounts have been reported to be expressed there (**Fig. S1 g**) (Lenis et al., 2018).

### Detergent free purification of ABCA4 with CL2 antibody and electron microscopy imaging

SMA-extracted bovine ROS was incubated with CL2-conjugated immunoaffinity resin (**Fig. 1 c**). Elution of ABCA4 with the known epitope peptide produced a concentrated and pure sample of ABCA4 (**Fig. 1 c**, “Elu” & “EW” lanes), with large amounts of elution peptide and characteristic SMA-smearing seen at the bottom of these lanes. The elution and subsequent wash from the immunoaffinity purification were then pooled and concentrated for size exclusion chromatography (SEC) (**Fig. 1 d**). To characterize possible morphological changes to ABCA4 in the SMALP, the purified samples were prepared for negative stain transmission electron microscopy (nsTEM). nsTEM micrographs showed monodisperse, homogenous ABCA4 particles (**Fig. S1 h**, left). Clear 2D class averages were made from particles picked by an unbiased autopicking feature of cisTEM (**Fig. S1 h**, right) (Grant et al., 2018). The resultant *de novo* 3D model, obtained using the cisTEM’s *de novo* reference map generator, showed significantly more density in the transmembrane domain (TMD) region than the prior nsTEM-generated structure of ABCA4 (**Fig. S1 i**). After refining, the roughly 4 nm-thick TMD showed a diameter of roughly 12 nm, which was larger than the previously published nsTEM model in the presence of detergent (EMDB-5497, orange) (**Fig. S1 j**) (Tsybovsky et al., 2013). The increased density did not confirm the presence of lipids in the TMD, and the possibility existed that more stain could have adhered to the TMD of the SMA-extracted protein. When considered in light of all of the results reported herein, however, we suspect the increased density was due to copurifed lipids. The other proportions obtained agreed well with the published ABCA4 nsTEM model and the general size and shape of ABCA1 (EMDB-6724, purple ribbon), determined by cryoTEM in 0.06% digitonin (**Fig. S1 i and j**) (Qian et al., 2017).

### Detergent free purification of PRPH2/ROM1 with novel nanobody Nb19

We developed a novel nanobody to pulldown PRPH2/ROM1 *via* an added His_6_ tag on the nanobody (**Fig. 1 e and f**, **Fig. 2**). All nanobodies share similar topology, they primarily vary in the hinge regions (H1, H2, etc.) which, upon folding, create complimentary determining regions (CDR’s) that constitute the paratope (**Fig. 2 a-c**). We selected, purified, and expressed 5 Nbs (Nb13, Nb19, Nb20, Nb28, and Nb32) representing different sequence families, each family grouped by CDR sequences (**Fig. 2 b and d**) (Pardon et al., 2014). All of the nanobodies bound tightly to pre-purified PRPH2/ROM1 sample as monitored by SEC (**Fig 2 e**). Nb19 proved to be the most efficient binder of the five; immunoprecipitation of PRPH2/ROM1 from extracted ROS (utilizing the His_6_ tag on the nanobody to bind Ni^2+^-resin) gave the highest yield (**Fig. 2 f**). The resulting PRPH2/ROM1-Nb19 complex was of sufficient purity after elution from Ni^2+^-resin to analyze its copurifying lipids directly (**Fig. 1 e**).

**Figure 2.**
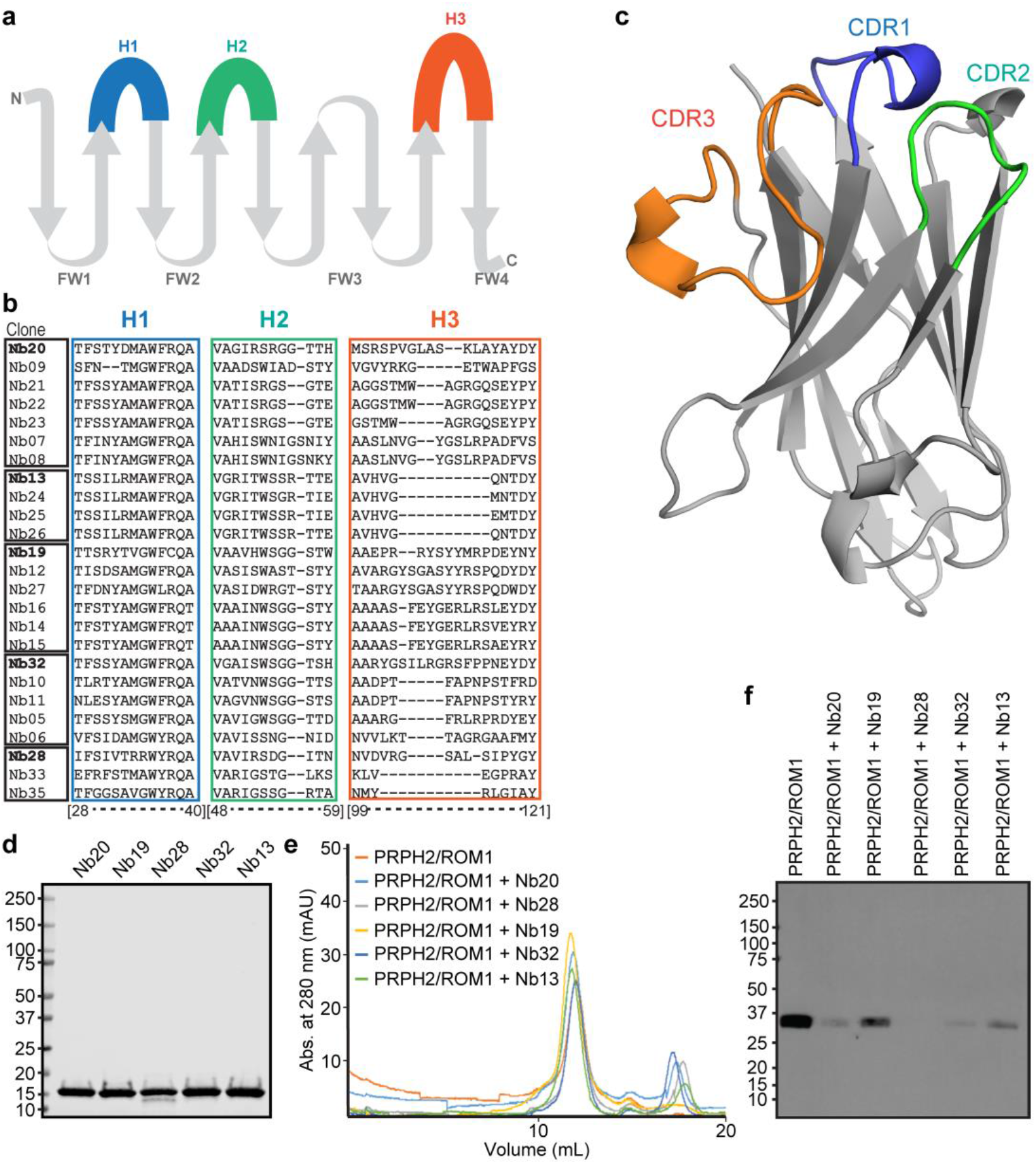
Biochemical characterization of the PRPH2/ROM1/Nb19 complex. (**a**) The secondary structure of the Nb domain consists of 9 beta sheets separated by loop regions. H1, H2, and H3 are separated by four framework regions (FWR’s). (**b**) Each of the five delineated Nb families are defined by boxes around the clone names. Hypervariable region sequences H1, H2, and H3 are listed after each clone name and boxed in blue, green and orange respectively. (**c**) Robetta-homology modeled Nb19 is shown, highlighting extended CDR regions encoded by hypervariable regions defined in (b). (**d**) 10 µg of purified Nb20, Nb19, Nb28, Nb32, and Nb13 were subjected to SDS-PAGE to indicate purity (stained with Coomassie Blue R250). (**e**) 10 µg of PRPH2/ROM1 was subjected to SP-200 gel filtration alone or after incubation with 20 µg of Nb. Nb19 caused the greatest shift in volume of elution. (**f**) Immunoprecipitation of PRPH2/ROM1 from solubilized rod OS with Nbs. First lane, purified PRPH2/ROM1 (1.0 µg), was used as a positive control. Detection of PRPH2/ROM1 was performed by immunoblotting with the C6 (anti-PRPH2) and 2H5 (anti ROM1) antibodies. Nb19-mediated immunoprecipitation produced the greatest quantity of PRPH2/ROM1.

### SMALP-encapsulated ABCA4 and rhodopsin retain ligand binding capacity

Assessing the activity of ABC transporters in SMA presents a challenge because the low millimolar concentrations of magnesium preferred for efficient coordination of ATP to the Walker A binding site of ABC transporters precipitates SMA (Oluwole et al., 2017). The correct folding and nucleotide binding of ABC transporters in SMALPs can be assessed *via* tryptophan fluorescence quenching with increasing concentrations of ATP in the absence of magnesium (Gulati et al., 2014). Using this assay, we confirmed that the SMA-purified ABCA4 is able to bind ATP (K_D_ = 133.5 µM), albeit with lower affinity than reported in the presence of magnesium (**Fig. 3 a**) (Ahn et al., 2000).

**Figure 3.**
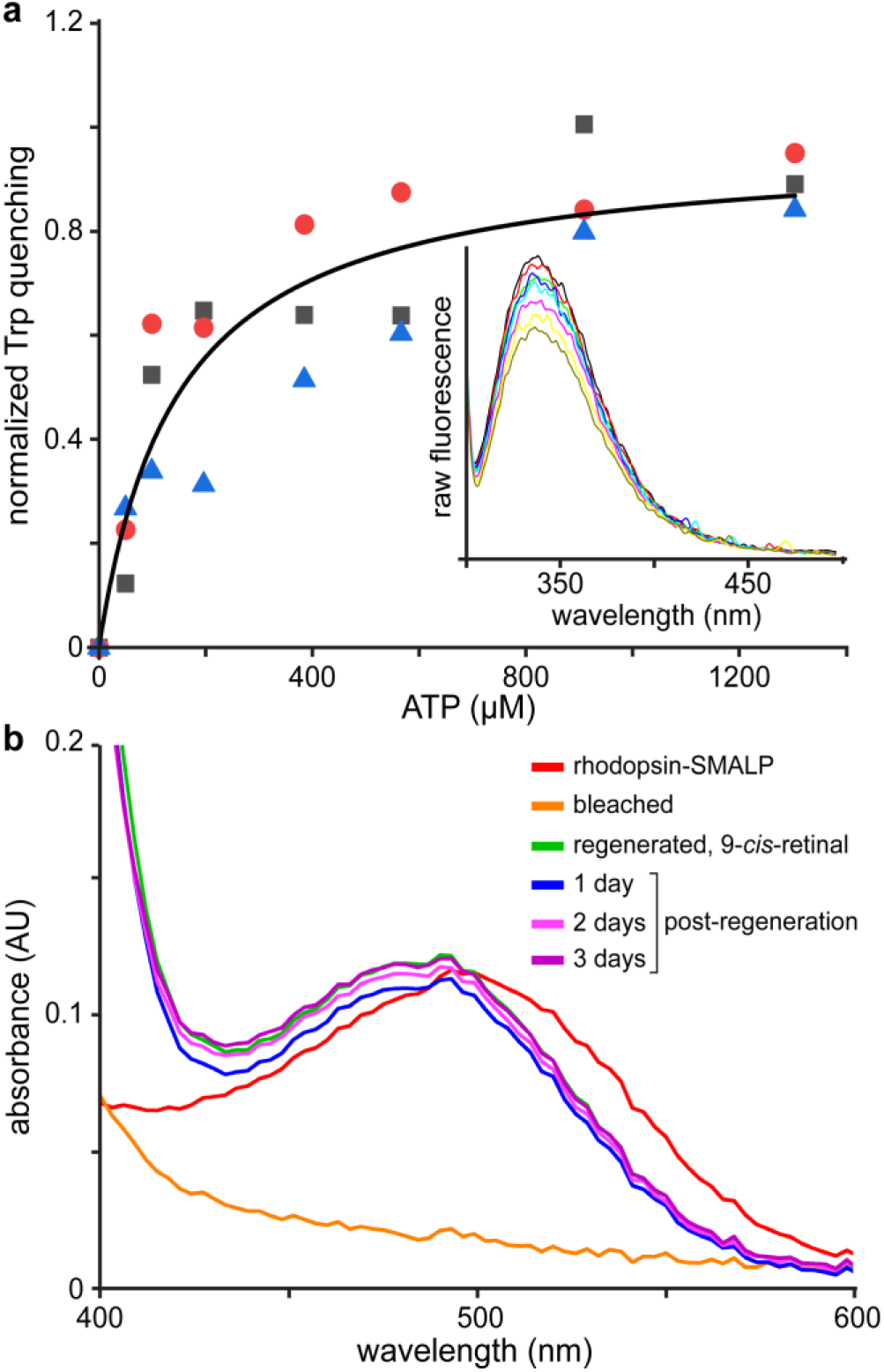
SMALP-encapsulated proteins retain ligand binding capacity. (**a**) ABCA4 extracted and purified in SMALPs shows intrinsic Trp-quenching characteristic of ATP transporters in the presence of serially-added ATP (K_D_ = 133.5 µM). Three separate experiments are shown with different symbols. Langmuir binding isotherm curve (black) fit to the average of 3 runs (B_max_ = 11.85%, 17.91%, and 10.00% for black, red, and blue, respectively). **Inset**: One set of spectra for increasing concentrations of ATP, showing diminution of raw fluorescence. (**b**) Absorption spectra of purified rhodopsin in SMALPs. Rhodopsin extracted and purified in SMA retains the chromophore throughout purification in the dark (red). Rhodopsin is able to be bleached when exposed to bright light and hydroxylamine and then regenerated by addition of 9-*cis*-retinal. The regenerated rhodopsin sample (Regen. 9-cis) retains the chromophore over several days at room temperature.

We also assessed the ability of rhodopsin to bleach and regenerate in SMALPs (**Fig. 3 b**). Rhodopsin was purified using the 1D4 antibody that had been developed previously and is well established for protein purification in detergent-solubilized conditions (**Fig. 1 g**) (Molday and Molday, 2014). The protein retained its chromophore when purified in the dark, which suggested that the rhodopsin was purified intact and could hold the chromophore while in the SMALPs. We subsequently found the SMA-purified rhodopsin was able to efficiently bleach by light, with and without hydroxylamine to scavenge the chromophore, showing that the protein either has sufficient free volume or the SMALP has enough flexibility to undergo conformational changes. Rhodopsin was also able to regenerate efficiently with 9-*cis*-retinal, as shown by the reappearance of the characteristic absorbance peak of the opsin-chromophore complex at 487 nm (Hubbard and Wald, 1952). The regenerated samples were stable and soluble for days at room temperature. These results highlight the ability of SMALPs to efficiently extract this model GPCR in a stable form from its native, mammalian tissue, as has been done in cell lines and lower species with other GPCRs (Jamshad et al., 2015; Gakhar et al., 2020; Ganapathy et al., 2020; Bada Juarez et al., 2020; Routledge et al., 2020; Ueta et al., 2020).

### Untargeted lipidomic analysis of native SMALPs documents different membrane environments for ABCA4, PRPH2/ROM1, and rhodopsin

With SMALP-extracted, immunopurified samples of these three representative membrane proteins in hand, we carried out a high-resolution study of the lipid environments of each protein. Lipidomic analysis indicated that the SMALPs were able to extractmore than phospholipids from native ROS membranes (**Figs. 4 and 5, Figs. S2-5**). We detected many metabolites and other lipids, including acylcarnitines (AcCa), ceramides (Cer), cholesterol esters (ChE), mono-di- and triacylglycerols (MG, DG, and TG, respectively), free fatty acids (FFA), cardiolipin (CL) and several lyso-phospholipids (LPC, LPE, and LPA). There were many distinctions in the relative species composition within these lipid classes. In general, we observed the samples of SMA-extracted ROS (starting material) and rhodopsin had similar compositions (as would be expected given the large share of the ROS occupied by rhodopsin). Likewise, we found that the SMALPs of ABCA4 and PRPH2/ROM1, which both reside in the rim region, had similar species distributions within each lipid class. As a percentage of the total lipid class, the samples from the rim region lacked AcCa(16:0), which was balanced by a relative enrichment of AcCa(22:4) (**Fig. 4 a**). There was no gradual increase in the chain lengths up to AcCa(22:4) in the rim samples, suggesting that free carnitine becomes conjugated to the 22:4 FA directly, and that the resultant AcCa(22:4) is not metabolized as quickly as species of similar length. The rim samples showed a relative abundance of Cer(d18:1_18:0) as compared to the rhodopsin samples, whereas the rhodopsin samples showed a relative abundance of Cer(d18:1_22:0), Cer(d18:1_24:1), and Cer(d18:2_24:0), suggesting a preference for longer chain lengths (**Fig. 4 b**). The same relative preferences were seen with LPC and LPE analyses. The rim samples showed significant enrichment in LPC(18:0) and LPE(18:0), while LPC(22:5) and LPE(22:5), as well as LPC(22:6) and LPE(22:6), were several-fold higher in the rhodopsin samples (**Fig. 4 c and d**). Cholesterol levels were found to be higher in rhodopsin samples when compared to the PRPH2/ROM1 samples, while ChE(18:2) was relatively enriched in the PRPH2/ROM1 samples relative to rhodopsin samples (**Fig. 4 e**).

**Figure 4.**
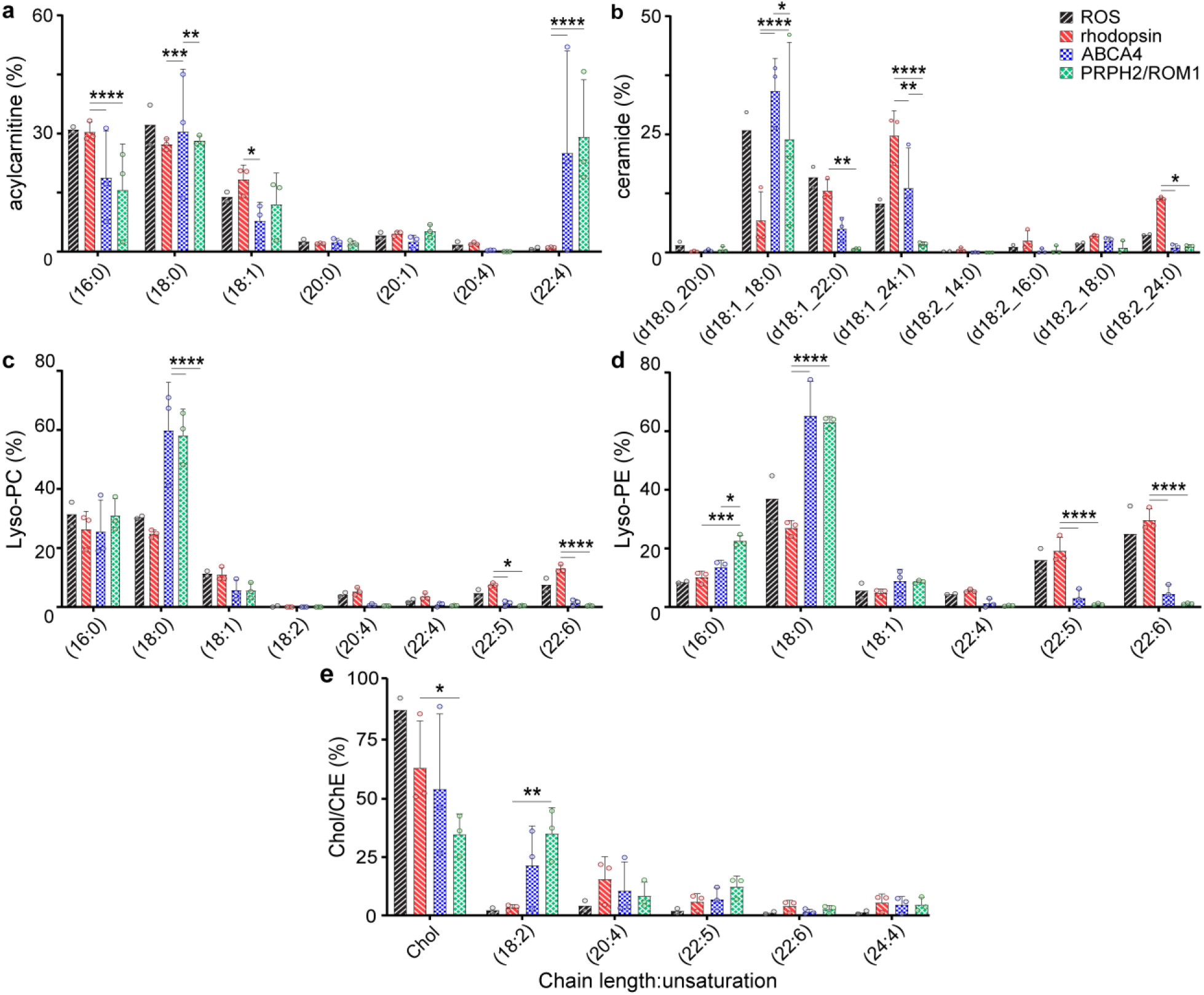
Lipid compositions of SMALP-embedded ROS membrane proteins are distinct to their native location in the membrane. **(a-e)** Percentages are shown of every detected species of AcCa, Cer, LPC, LPE, and cholesterol/ChE, respectively, extracted from SMALPs. Selected species are graphed (all species are shown in **Figs. S2-5**). Total ROS: black forward stripe; rhodopsin: red backward stripe; ABCA4: blue checker; PRPH2/ROM1: green diamond. ROS measured in duplicate as noted by individual data points (open circles). Percent composition was derived from each sample by dividing the area under the curve for each species in a class by the total area under the curve for the class reported *via* LC-MS after correction for variations in internal standard area, sample mass, and sample injection volume. Statistics were determined using two-way ANOVA with Tukey’s multiple comparisons post-hoc test. Significance values are indicated as follows: *, *P* < 0.05; **, *P* < 0.01; ***, *P* < 0.001; ****, *P* < 0.0001.

**Figure 5.**
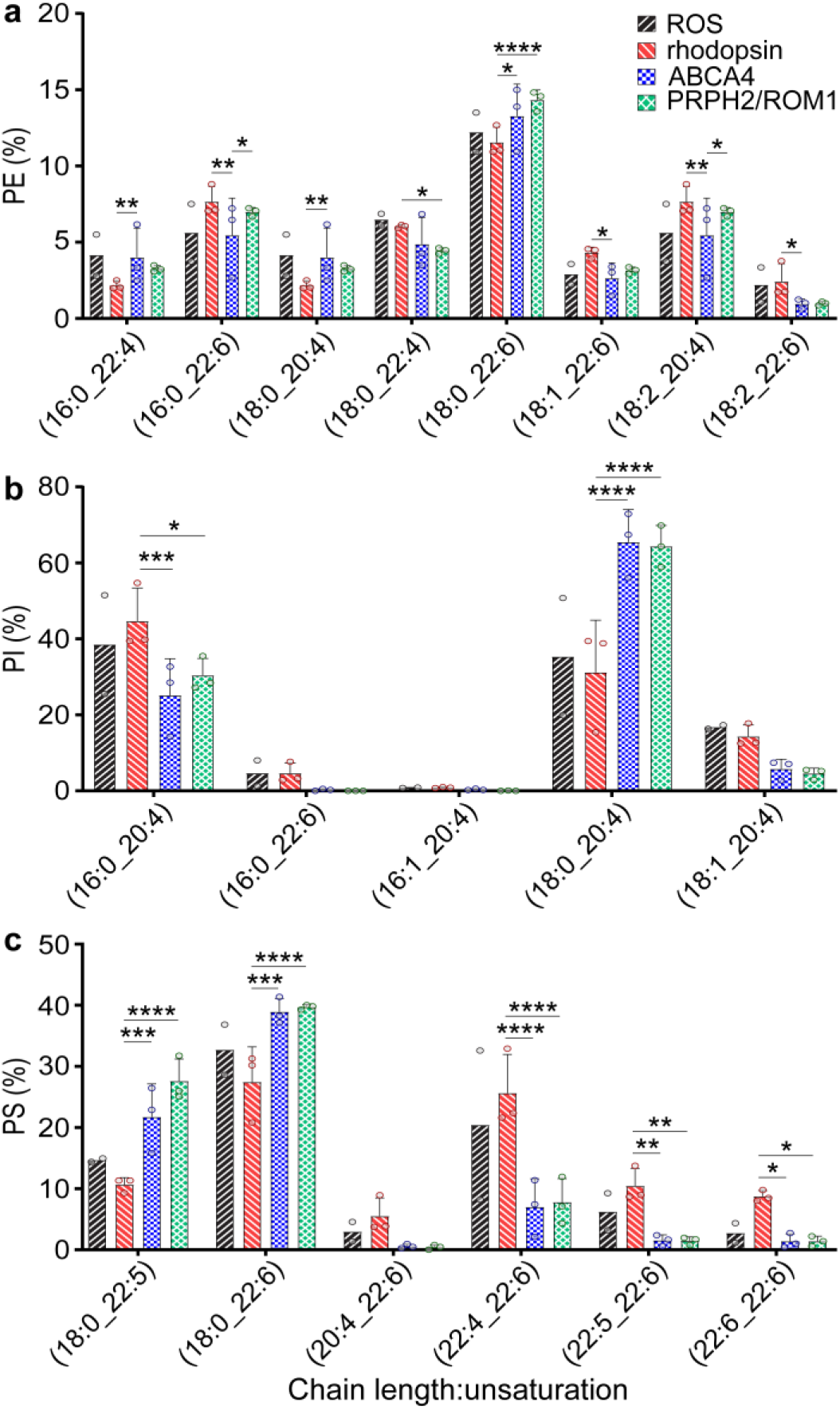
Phospholipid compositions of SMALP-embedded ROS membrane proteins are distinct to their native location in the membrane. **(a-c)** Percentages are shown of every detected species of PE, PI, and PS, extracted from SMALPs; selected PL species are shown here (all PL species are shown in **Figs. S2-5**). Total ROS: black forward stripe; rhodopsin: red backward stripe; ABCA4: blue checker; PRPH2/ROM1: green diamond. ROS measured in duplicate as noted by individual data points (open circles). Major differences are evident between ABCA4 and PRPH2/ROM1 (rim) and rhodopsin (center). Percent composition was derived for each sample by dividing the area under curve for each species in a class by the total area under curve for the class reported *via* LC-MS after internal standard, sample mass, and sample injection volume correction. Statistics were determined using two-way ANOVA with Tukey’s multiple comparisons post-hoc test. Statistical significance values are indicated as follows: *, *P* < 0.05; **, *P* < 0.01; ***, *P* < 0.001; ****, *P* < 0.0001.

The common phospholipids also displayed multiple significant differences between the rim region and the center (**Table 1**), especially between PC and PE. There were many differences at the species level within each PL class as well (**Fig. 5**). There were some instances of differences in PE species between the samples of the two rim proteins, where rhodopsin and PRPH2/ROM1 were relatively higher in PE(16:0_22:6) and PE(18:2_22:6) when compared to ABCA4 (**Fig 5 a**). There were also significant differences among individual species in the phosphatidylinositol (PI) and PS classes (**Fig. 5 b and c**). Here, though, the rim samples ha similar profiles and were both distinct from rhodopsin samples, further confirming the similarity between the rim sample membranes.

**Table 1.**
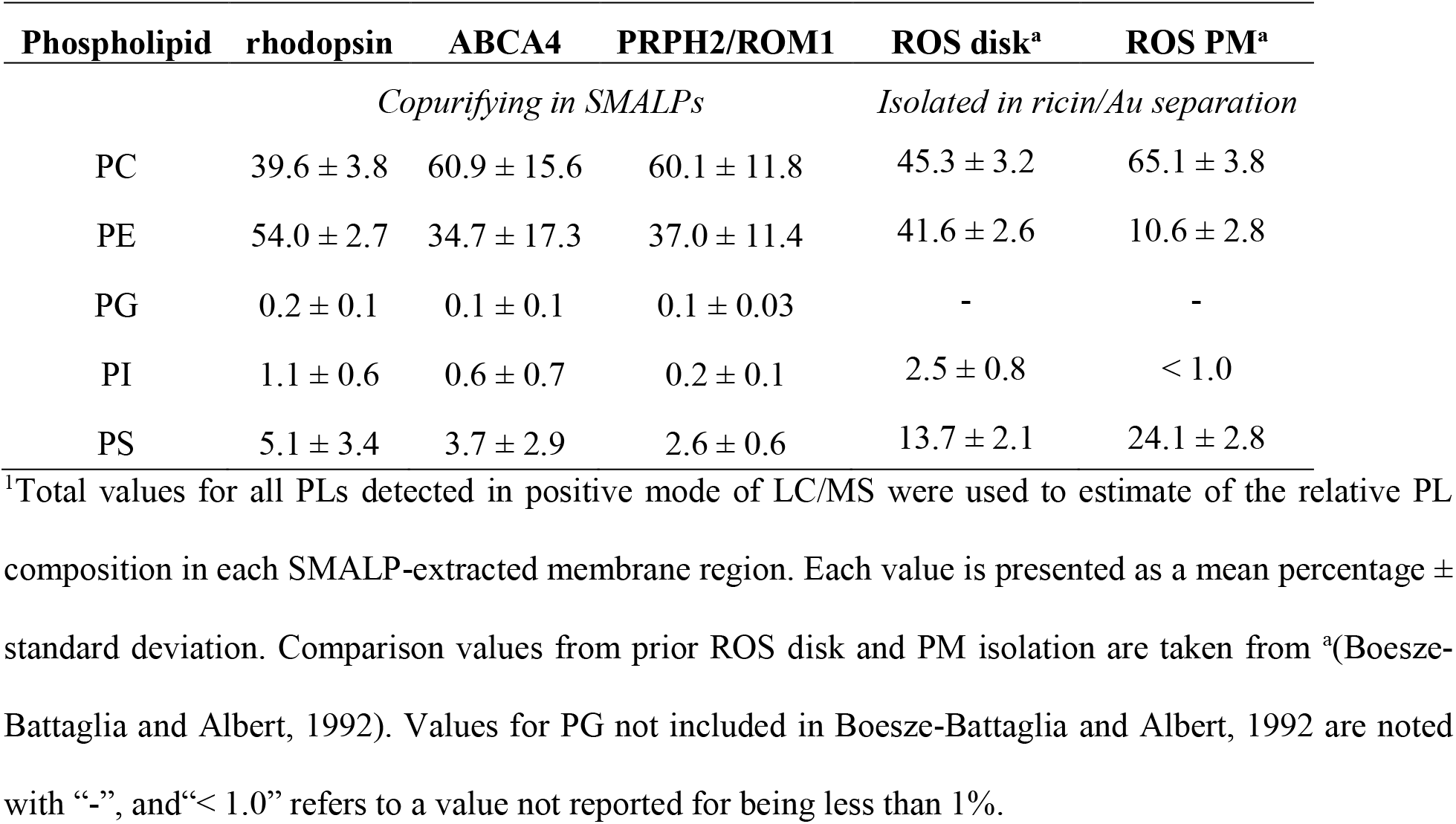
Comparison of relative PL compositions in native ROS membrane domains.

We further evaluated the aggregate relationship between each sample using the unbiased method of principle component analysis (PCA) (**Fig. 6**). PCA was able to simplify the relationships between 199 separate species across 14 lipid classes and found similarity between the rim samples and difference between the rim and the center region samples. Principle components 1 and 2 (PC1 and PC2, respectively) totaled a combined 64.16% of the variance in the system, with PC1 accounting for over 46%. The resultant PCA scores showed clustering of ABCA4 and PRPH2/ROM1 samples along both PC1 and PC2, far removed from rhodopsin along the PC1 axis. The rhodopsin samples grouped tightly, and associated more closely with the starting ROS samples with respect to PC1. Analysis of the PCA loadings suggested that PC1 found strongest differences in species across classes containing palmitic and stearic acid (16:0 and 18:0, respectively) (corresponding to the rim samples) and chain lengths 20 or more containing 4-6 unsaturated bonds (rhodopsin samples). We conclude that the lipid composition of the rim and center regions of ROS disks are distinct at the lipid species level.

**Figure 6.**
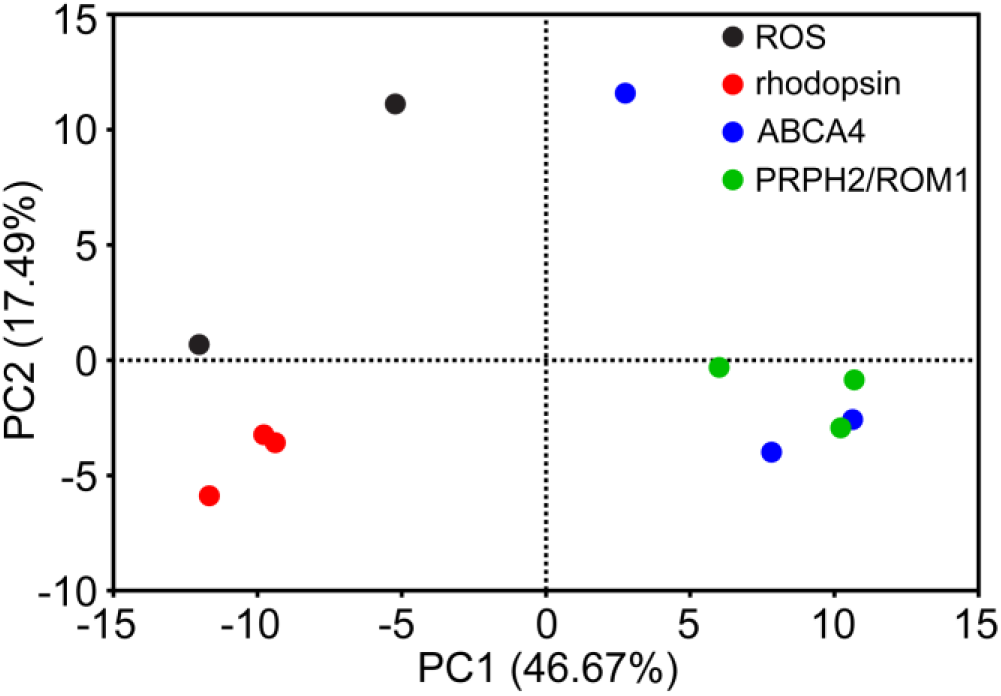
PCA groups rim region samples apart from central, rhodopsin samples. PCA of 199 lipid species from 14 lipid classes shows clustering of rim samples away from rhodopsin samples, highlighting, in an unbiased manner, the similarity of the rim membrane lipids and the center region lipids. PC1 and PC2 combined to equal 64.16% of the total variance. PC1-3 had eigenvalues greater than the 95^th^ percentile of eigenvalues randomly generated through parallel analysis (1000 simulations conducted), but PC3 was not used because of its lower proportion of the variance (13.51%).

### Comparisons between the central and rim regions of ROS disks show differences in FA composition

The PCA results suggested that FA chain length and/or unsaturation of the lipids residing in these two functionally distinct areas may be a key differentiator between their membranes. To address this fully, we performed lipid extractions from each SMALP-protein sample, then hydrolyzed the head groups of all lipid species in each sample, followed by FA lipidomic analysis *via* LC/MS. The FA compositions of the lipids isolated from the two rim-region proteins ABCA4 and PRPH2/ROM1 showed no statistically significant differences in relative molar percent for all chain lengths and saturations. There was considerable difference, however, in the FA composition of the rhodopsin-containing samples when compared to the rim proteins. The rim region proteins copurified with predominantly unsaturated and short chain length FAs, especially 16:0 and 18:0 (**Fig. 7 a**). Those two FA species accounted for over 67% relative to the entire FA content of the ABCA4 sample; and over 82% of the PRPH2/ROM1 sample. Conversely, the rhodopsin samples contained less than 30% of these two FAs.

**Figure 7.**
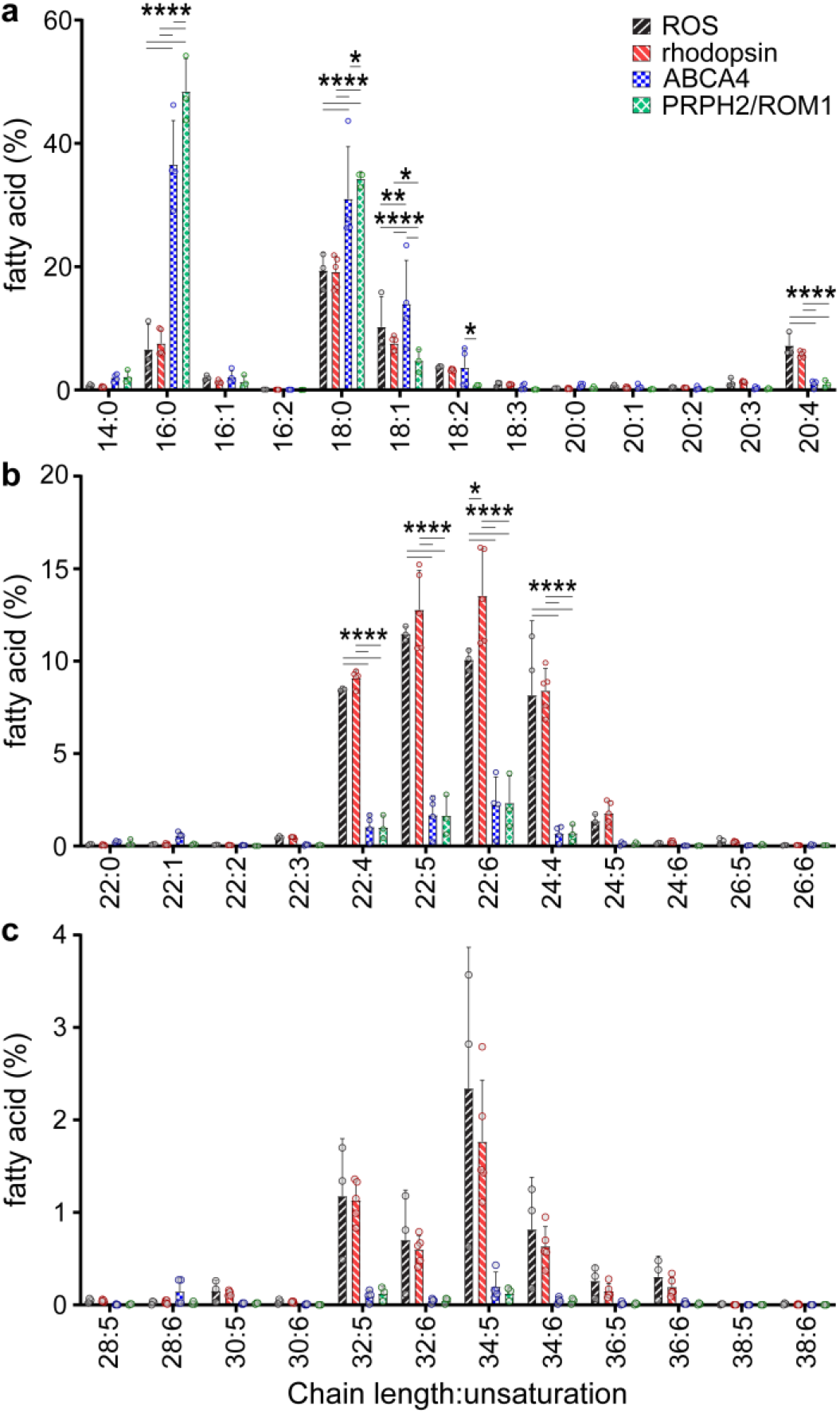
Comparison of FA chain lengths between the center and rim of ROS disks shows relative enrichment of shorter chain lengths in the rim and LC- and VLC-PUFAs in the center. (**a**) Relative molar percentages are shown of every detected class of FA molecule (C14-20) extracted from the SMALPs of each purified protein. (**b**) Relative molar percentages are shown of every detected class of FA molecule (C22-26) extracted from the SMALPs of each purified protein. (**c**) Relative molar percentages are shown of every detected class of FA molecule (C28-38) extracted from the SMALPs of each purified protein. Total ROS: black forward stripe; rhodopsin: red backward stripe; ABCA4: blue checker; PRPH2/ROM1: green diamond. The significance of differences between the means was determined using two-way ANOVA with Tukey’s multiple comparisons post-hoc test. ROS: n = 3, rhodopsin: n = 5, ABCA4: n = 4, PRPH2/ROM1: n = 3. Significance values are indicated as follows: *, *P* < 0.05; **, *P* < 0.01; ***, *P* < 0.001; ****, *P* < 0.0001.

Docosahexanoic acid (22:6, DHA) is known to be essential to ROS disk health, and DHA has been shown to facilitate rhodopsin activity (Bush et al., 1991; Organisciak et al., 1996; Litman et al., 2001). We found DHA was significantly higher in the central region than in the rim, with a DHA relative molar percent of 13.5% for rhodopsin samples (**Fig. 7 b**). The rhodopsin samples were enriched in LC-PUFAs more generally as well, whereas the rim samples contained only 1.6% or less molar percent LC-PUFAs.

Rhodopsin SMALPs also contained more VLC-PUFAs than those in the disk rim (**Fig. 7 c**). The most prominent VLC-PUFAs found in rhodopsin samples were dotriacontapentaenoic, dotriacontahexaenoic, tetratriacontapentaenoic, and tetratriacontahexaenoic acid (32:5, 32:6, 34:5, 34:6, respectively), with relative abundances between 0.6% - 1.3%. In contrast, the rim protein SMALPs were sparsely populated with VLC-PUFAs, accounting for 0.2% or less of their total FA content.

## DISCUSSION

The first question to be answered by this study is whether lipids that copurify in SMALPs containing purified membrane proteins faithfully represent the native membrane regions from which they are purified. There have been reports that SMALPs composed of pure phospholipids of different types (*e.g*., PC vs PE) rapidly exchange when incubated together, suggesting that native tissues, left solubilizing in SMA for 1-2 h, would result in an equilibrated distribution of membrane components among the protein-containing SMALPs (Cuevas Arenas et al., 2017; Schmidt and Sturgis, 2018; Danielczak and Keller, 2018). Prior work on single target proteins purified in SMALPs from membranes showed little difference between the mother membrane and the extracted, copurifying lipids (Dörr et al., 2014). Here, we document definitive differences between samples isolated from different regions of the same mammalian membrane tissue, analogous to a recent report on bacterial membrane proteins (Teo et al., 2019). One explanation for the lack of predicted homogeneous mixing could be that the diverse membrane constituents of native membranes are organized into protein-dependent subdomains that are not susceptible to the fast exchange seen with pure phospholipid nanodiscs. Nevertheless, spontaneous lipid exchange is known to occur in biological membranes, so there is evidence in nature for membrane components to swap particular lipids in a manner that approaches equilibration (Bell, 1978). The bounded, roughly 12 nm-diameter SMALP disks may not behave like biological membranes, so ambiguity remained as to whether the lipid compositions in our study were the result of preferential sorting after extraction, or in fact represented the hyper-local membrane environments of the purified proteins.

To address the possibility of preferential sorting after extraction, we purified two proteins from the same rim region, ABCA4 and PRPH2/ROM1, and compared their lipid profiles with that of rhodopsin. We hypothesized that if the local lipid environment is preserved in SMALPs, then samples of the rim-region proteins should show similar lipid profiles to one another, distinct from that of rhodopsin samples. Our FA chain length/unsaturation analysis revealed no statistically significant differences between the two rim region samples; and indeed, there were clear differences between the FA arrays of the rhodopsin and rim samples. Furthermore, the cases of statistically significant differences between each rim sample and rhodopsin were nearly identical across all FA chain lengths and saturation levels. Therefore, this study provides strong evidence that SMA-extracted samples from native tissue are highly likely to retain the local environment from which they were isolated.

Rhodopsin is known to achieve a paracrystalline state in the central, flat portion of ROS disks, but it has also been found in the plasma membrane (Fotiadis et al., 2004; Kessler et al., 2014). We estimate that rhodopsin in the plasma membrane should account for less than 2% of total rhodopsin purified herein, given the calculated amount of rhodopsin in the plasma membrane of murine ROS is 2% (Kessler et al., 2014). The increased size of bovine ROS should decrease the relative amount of rhodopsin in the plasma membrane as the disk membranes scale in cumulative surface area more quickly than the plasma membrane as dimensions increase. ABCA4 and PRPH2/ROM1, conversely, have been shown to localize on the rim region of the disks and have not been shown to exist in the plasma membrane in detectable amounts (Molday et al., 1987; Illing et al., 1997).

We approximated the amount of the ROS disk membrane accounted for by the SMALPs of our three chosen samples to assess the completeness of our analysis. We estimate 95% of the disk membranes are accounted for in the combined SMALPs of rhodopsin, ABCA4, and PRPH2/ROM1 (**Table 2**). We arrived at this estimation by finding the proportion of rhodopsin, ABCA4, PRPH2, and ROM1 in comparison with the other membrane components of ROS and comparing their theoretical numbers of copurifying lipids to get a weighted lipid contribution (WLC) for each protein (**Eq. 1**).

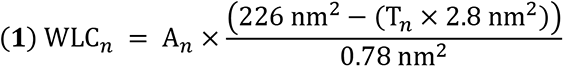

**Table 2.** Comparison of weighted lipid composition of ROS disk-specific membrane proteins.

| Disk Protein | APEX<sup>a</sup> | A<sub>n</sub> | T<sub>n</sub> | # lipids per protein | WLC | WLC (%) |
| --- | --- | --- | --- | --- | --- | --- |
| rhodopsin | 0.1580841 | 0.663 | 7 | 264.6 | 175.3 | 68.3 |
| PRPH2/ROM1 | 0.0614938 | 0.258 | 16 | 232.3 | 59.9 | 23.3 |
| ABCA4 | 0.0073837 | 0.031 | 12 | 246.7 | 7.6 | 3.0 |
| GC-1 | 0.0073970 | 0.031 | 1 | 286.2 | 8.9 | 3.5 |
| R9AP | 0.0016516 | 0.007 | 1 | 286.2 | 2.0 | 0.8 |
| ATP8A2 | 0.0013005 | 0.005 | 10 | 253.8 | 1.4 | 0.5 |
| GC-2 | 0.0012608 | 0.005 | 1 | 286.2 | 1.5 | 0.6 |

<sup>1</sup>Absolute protein expression (APEX) levels are taken from <sup>a</sup>(Kwok et al., 2008). A<sub>n</sub> is each the APEX value of each protein, n, divided by the sum of all APEX values of disk-specific proteins. T<sub>n</sub> is the number of transmembrane helices of each disk-specific protein, n. WLC is the calculated weighted lipid composition of the theoretical SMALP of each protein, as described by **Eq. 1**. GC-1 and -2 are guanylyl cyclase 1 and 2, respectively. R9AP is regulator of G protein signaling 9-binding protein. ATP8A2 is ATPase
aminophospholipid transporter type 8A, member 2. Based on the estimation of WLC percent in the last column, rhodopsin, PRPH2/ROM1 and ABCA4 SMALPs account for 95% of the membrane lipids of ROS disks when extracted in SMA.

We cross-referenced the nine membrane proteins classified as ROS disk-specific by Skiba et al. with the ROS disk proteomics reported by Kwok et al. using absolute protein expression (APEX) measurements taken by MS/MS (Skiba et al., 2013; Kwok et al., 2008). Each protein was normalized to their proportion of disk-specific membrane protein by dividing its APEX amount by the APEX amount for all disk-specific proteins combined (A_n_, where n is a disk-specific protein, **Eq. 1**). These percentages were used to weight the theoretical number of lipids per SMALP for each protein. The theoretical number of lipids per SMALP for each protein was determined by subtracting the product of each protein’s number of TM helices (T_n_, **Eq. 1**) and twice the average cross-sectional area of a transmembrane alpha helix (∼1.4 nm^2^) from the theoretical total surface area of a 12 nm diameter SMALP (∼113 nm^2^ per side of disk) (**Fig. S1 i**) (Eskandari et al., 1998; Swainsbury et al., 2014; Takamori et al., 2006). This number gave the free area for lipids in each SMALP, and an estimate of the number of lipids was calculated by dividing by the average cross-sectional area of a phospholipid (∼0.78 nm^2^) (Lee, 2003). Each of the disk-specific membrane proteins’ average number of phospholipids per SMALP was then weighted by A_n_, resulting in the WLC of each protein. WLC_rhodopsin_, WLC_ABCA4_, and WLC_PRPH2/ROM1_ were added together and divided by the sum of all WLCs, giving an approximate lipid contribution of 95% from the three samples studied here (**Eq. 2**).

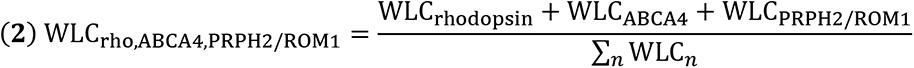

This estimation gives us confidence that we have studied the majority of the disk membranes.

The stark contrast in the profiles of FA chain lengths between the rim and center of the disks is remarkable (**Fig. 7**). The center of the disk is enriched with LC- and VLC-PUFAs relative to the disk rim. The relative abundance of DHA coincident with rhodopsin is consistent with the well-documented requirement of DHA for healthy rhodopsin activity (Mitchell et al., 1992b). The relative abundance of eicosatetraenoyl acid (arachidonic acid (AA), 20:4) in the disk center is consistent with its well-known role as a critical precursor for LC- and VLC-PUFAs (Grogan and Lam, 1982; Grogan and Huth, 1983; Grogan, 1984).

On a more general scale, the rhodopsin samples showed that combined VLC-PUFAs represent over 15% of total FAs in the center of bovine ROS disks, roughly equivalent to the 13% of whole bovine ROS reported by Aveldaño and Sprecher (using their classification of VLC-PUFA as ≥24 carbons in length) (Aveldaño and Sprecher, 1987). A particularly intriguing finding was the distinct lack of VLC-PUFAs in the rim region. We had surmised that the slightly wider and curving rim region might provide more space for the extended acyl chains of VLC-PUFAs, but we now deduce that the rim region membranes require the stiffness provided by the abundant 16:0 and 18:0 saturated chains found there.

We were struck by the panoply of components extracted by SMALPs and measured by the lipidomic analysis of the ROS. Many membrane components copurified with the sample proteins, including lyso-PLs, sterols, sphingolipids, AcCa, FFA, cardiolipin, and mono/di/triglycerides. This comprehensive report **(Figs. 4 and 5, Fig. S2-5)** of the components of the ROS disk membranes is, to our knowledge, the most complete of any tissue extracted by native nanodiscs (*e.g*., SMALPs). The results of our PCA confirm, in an unbiased manner, that many of the diverse components found in this study are spread anisometrically across the continuous ROS disk membrane, favoring the center or rim region (**Fig 6**). Some of this systematic heterogeneity is likely critical to the maintenance of healthy phototransduction and should be probed more deeply. This data also begs the question of how the asymmetry is initiated and maintained by ROS membrane proteins.

Differences in content of acyl-carnitine and lyso-PLs are the most notable in the analysis of the lipid classes. While carnitine has been reported to be in ocular tissues, our data further localize at least some of the acyl-carnitine to the center of the ROS disks (**Fig. 4 a**) (Pessotto et al., 1994). Previous work has shown that injection of carnitine in the eye can be protective in a methylcellulose-induced ocular hypertension model, as measured by decreased levels of inducible nitrogen oxide synthase (iNOS), malondialdehyde (MDA), and ubiquitin (Ub) (Calandrella et al., 2010). Our results, which place AcCa in the immediate vicinity of rhodopsin in the membrane, suggest that carnitine may act as a check on normal oxidative stress in the OS disk membranes. Supplemental carnitine could increase the protective effect afforded the retina by endogenous levels of carnitine in the OS disks, but more evidence is needed to confirm this.

The presence of lyso-PLs has been reported at the tissue level in bovine and human retinas, but their specific function(s) in the retina are yet to be determined (Berdeaux et al., 2010). As a surfactant, lyso-PC has been shown to increase membrane fluidity, which is itself important for protein reorganization in OS disks (Henriksen et al., 2010; Rakshit et al., 2017). Our data show unequivocal differentiation of the lyso-PLs, with short, saturated species on the rim and LC-PUFAs in the center (**Fig. 4 c and d**). Lyso-PLs consisting of LC-PUFAs likely contribute even more fluidity to the center of the disk. In addition to this general effect, it is conceivable that the lyso-PLs in the center of ROS disks interact specifically with the membrane proteins in a signaling capacity. Lyso-PLs have been shown to interact with GPCRs to initiate G_12/13_, G_q/11_, G_i_ and G_s_ signaling, thereby affecting various downstream, intracellular signaling pathways (Anliker and Chun, 2004; Xiang et al., 2013; Torkhovskaya et al., 2007; Li et al., 2016). Rhodopsin is already known to be affected by the membrane composition when transitioning between the Meta I and Meta II states (Gibson and Brown, 1991a; b, 1993; Botelho et al., 2002), but more study is needed to probe the possibility of alternative G protein interactions with rhodopsin for the propagation of lyso-PL signals.

The trend indicating enrichment of free cholesterol toward the center of the disks was surprising (**Fig. 4 e**). Past theories suggested that an exchange of disk cholesterol with the PM causes a gradient of cholesterol from high (nascent disks) to low (mature disks). Therefore, we had expected to see a relative increase in free cholesterol in the rim of the disk (Boesze-Battaglia et al., 1989). One way to explain our result is that the rim, with its highly curved structure, cannot maintain high levels of cholesterol. There may be a separate route for cholesterol movement between disks that allows for the diminution of free cholesterol in maturing disks, but this is only speculation. Regardless, all samples isolated from the disks showed lower relative levels of free cholesterol than the ROS starting material, which contained both disks and PM.

This study, to our knowledge, is the first to extract and purify mammalian membrane proteins along with their corresponding native membrane environment. We were able to document the precise, species-level differences between the two lipid domains of ROS disks (**Fig. 8**). Our results could provide more context for prior work done on detergent-resistant membranes (DRMs) of the ROS, where Triton X-100-resistant membranes low in rhodopsin and seemingly high in ABCA4 were isolated from the rest of ROS disks (Martin et al., 2005). The DRMs were shown to have some of the same trends between DRM and fully-solubilized regions as seen between the rim and center regions in this work.

**Figure 8.**
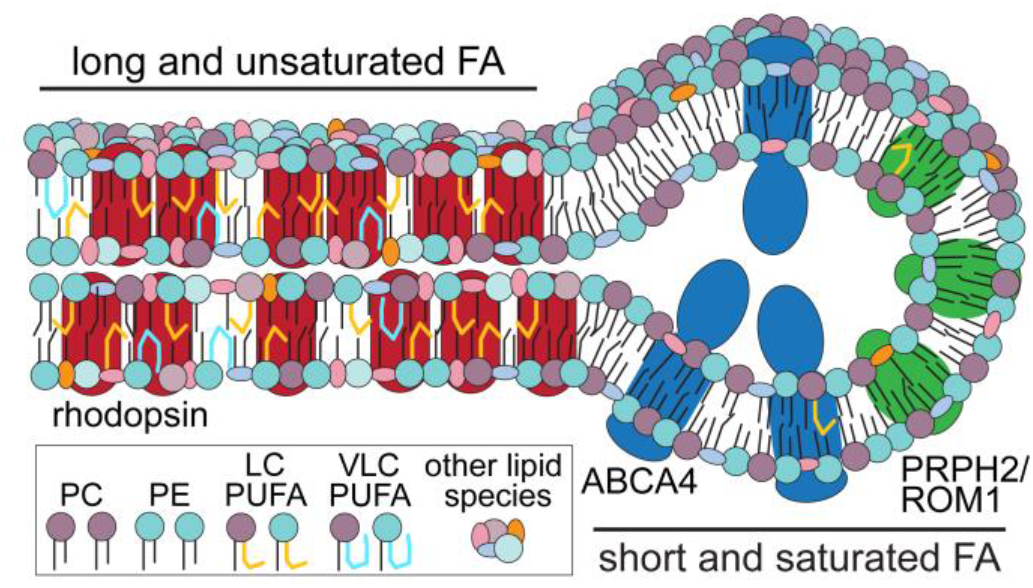
ROS disks have regionally distinct microenvironments. The center region of ROS disks, rich in rhodopsin, have an abundance of long and unsaturated FAs. Rim regions of ROS disks have relatively high amounts of short and saturated FAs. There are many other distinctions in lipid species between the two regions, including relative amounts of PC and PE.

Further work should be dedicated to studying physiological protein-lipid interactions of the retina, as many of the key proteins in the visual cycle and phototransduction are membrane proteins. To this end, the process of studying differential membrane composition based on native protein isolation in SMALPs should be expanded to other systems in the hope of uncovering detailed information on the preferred lipid environment of other membrane proteins. In particular, the use of high-resolution lipidomics may help explain pathologies involving critical protein-lipid interactions.

## EXPERIMENTAL PROCEDURES

### Animals

All animal protocols were approved by the Institutional Animal Care and Use Committees at the University of California, Irvine and were conducted in accordance with the Association for Research in Vision and Ophthalmology (ARVO) *Statement for the Use of Animals in Ophthalmic and Visual Research*. Wild-type (WT) and *Abca4*^-/-^*Rdh8*^-/-^ mice on a BALB/cJ background were used in this study. All mice were housed in the University Laboratory Animal Resources (ULAR) facilities at the University of California, Irvine and maintained in a 12 h/12 h light-dark cycle, and fed Teklad global soy protein-free extruded rodent diet (Envigo, Indianapolis, IN) chow and water *ad libitum*.

### Extraction of ROS proteins in SMA

Detergent (laurel maltose neopentyl glycol (LMNG)) (Anatrace, Maumee, OH) or XIRAN SL30010 P20 (Polyscope Polymers B.V., Netherlands) SMA (2.3:1 styrene:maleic acid ratio) were incubated at varying concentrations for 1 h with ROS obtained from 3-4 bovine retinas in 1 mL of Extraction Buffer (20 mM BTP, pH 7.9, 10% glycerol, 300 mM NaCl, 1 mM TCEP). The incubations with SMA were conducted at RT, and with detergent at 4 °C. All samples were centrifuged at 100,000*g* for 1 h, and the soluble fractions were separated. Each pellet was resuspended in 10% SDS-containing Wash Buffer. Ten µl were loaded for each sample onto a Mini-PROTEAN TGX precast gel, 4-20% gradient (Bio-Rad, Hercules, California), and in the case of immunoblot analysis, proteins were transferred to a PVDF membrane. After blocking for 1 h in 5% (w/v) non-fat dry milk, anti-ABCA4 primary antibody TMR4 was added at 1:1,000 dilution from a 1 mg/mL stock, and incubated with the membrane overnight at 4 °C. Membranes were washed with PBST and then anti-mouse IgG (H&L) alkaline phosphatase-conjugated secondary antibody (Promega) was incubated with the membrane at a 1:5,000 dilution for 1 h at RT. After the membranes were again washed with PBST, the blots were developed with Western Blue® Stabilized Substrate for Alkaline Phosphatase (Promega, Madison, WI) for roughly 15 sec, then quenched with ultrapure water.

### Immunoblotting of bovine ABCA4

ROS of 50 bovine retinas were isolated as described previously and suspended in Extraction Buffer (20 mM BTP, pH 7.9, 10% glycerol, 300 mM NaCl, 1 mM TCEP) containing 2% LMNG (Anatrace) (Papermaster, 1982). The soluble fraction was separated from insoluble material by centrifugation at 100,000*g* for 1 h at 4 °C. A 10 μL aliquot of the soluble fraction was loaded into each lane of a Mini-PROTEAN TGX precast gel, 4-20% gradient (Bio-Rad,); and then proteins were transferred to PVDF membranes. After blocking for 1 h in 5% (w/v) non-fat dry milk, primary antibodies against ABCA4, namely CL2, Rim3F4, and TMR4 (Zhang et al., 2015) were added at dilutions of 1:10,000 from 1 mg/mL stocks, and incubated overnight at 4 °C. Membranes were washed with PBS containing 0.1% (v/v) Tween 20 (PBST), and then anti-mouse IgG (H&L) alkaline phosphatase-conjugated secondary antibodies (Promega) were incubated with the blots at a dilution of 1:5,000 for 1 h at room temperature (RT). After washing with PBST, blots were developed with Western Blue Stabilized Substrate for Alkaline Phosphatase (Promega) and imaged using an Odyssey Fc imager (LI-COR, Lincoln, NE), using the 700 nm channel with a 2-min exposure time.

### Immunoblot of Murine Retinas

Murine samples were obtained from the enucleated eyes of WT and *Abca4*^-/-^*Rdh8*^-/-^ mice according to a previously published protocol (Wei et al., 2016). Protein concentrations were determined with a BCA Assay kit (Bio-Rad), following the manufacturer’s instructions. Protein samples were mixed with NuPAGE LDS sample buffer and NuPAGE reducing agent, separated using NuPAGE 4-12% Bis-Tris gels (Invitrogen, Carlsbad, CA), and transferred to PVDF membranes. Membranes were blocked with 5% (w/v) non-fat dry milk and incubated with the CL2 antibody overnight at 4 °C. After washing with PBST, membranes were incubated with peroxidase-linked anti-mouse or anti-rabbit IgG (1:10,000) (Jackson ImmunoResearch Laboratories, West Grove, PA) for 1 h at room temperature. Protein bands were visualized after exposure to SuperSignal West Pico Chemiluminescent substrate (ThermoFisher Scientific, Waltham, MA).

### Immunohistochemistry of retinal sections

Mouse eye cups were fixed for 1 h in PBS containing 4% (w/v) paraformaldehyde (Sigma-Aldrich) at room temperature. After fixation, the eye cups were incubated sequentially in PBS containing 10, 20 or 30% (w/v) sucrose (Sigma-Aldrich, St. Louis, MO) for 30 min at room temperature. Then, the eye cups were infiltrated with a 2:1 mixture of PBS containing 30% sucrose and OCT compound (VWR International, Radnor, PA) and frozen with dry ice. Retinal sections were cut at a thickness of 12 μm and stored at −80 °C until use. The retinal sections were rehydrated with PBS and blocked with PBS containing 5% (v/v) goat serum (Thermo Fisher Scientific) and 0.1% (v/v) TritonX-100 (Sigma-Aldrich). After blocking, the sections were incubated with the appropriate primary antibodies diluted in PBS containing 5% goat serum overnight at 4 °C. Primary antibodies used for immunohistochemistry were Rim3F4, TMR4, and CL2. The retinal sections were washed with PBS three times for 5 min each and then incubated with Alexa Fluor 488-conjugated goat anti-mouse immunoglobulin G (IgG) diluted in PBS containing 5% goat serum at 1:400. After incubation, the retinal sections were washed with PBS three times for 5 min each and then mounted with Vectashield Mounting Medium (Vector Laboratories, Burlingame, CA). The images were acquired with a BZ-X810 Keyence microscope (Keyence, Itasca, IL) at 20X with numercal aperature of 0.75 at RT with no imaging medium and Alexa Fluor 488 used as the fluorochrome. The camera was built into the BZ-X810 Keyence microscope, and BZ-X800 viewer from Keyence was the acquisition software. Adobe Photoshop was used to adjust the orientations and Adobe Illustrator to make the figure.

### Purification of native, bovine ABCA4 in SMA

ROS isolated from 50 bovine retinas were extracted in 16 mL of Extraction Buffer with 2.5% SMA (v/v) (XIRAN SL30010 P20) (Polyscope Polymers B.V.) for 1 h at 4 °C in the dark, followed by centrifugation at 100,000*g* for 1 h at 4 °C. 1 mL of ∼8.0 mg/mL fresh immunoaffinity resin was prepared by conjugating purified, anti-ABCA4 antibody (CL2) to CNBr-activated Sepharose 4B beads (GE Healthcare Bio-Sciences, Chicago, IL, USA) according to manufacturer’s instructions. The extracted fraction of ROS was then mixed with the immunoaffinity resin, brought to 168 mM NaCl through dilution with SMA Wash Buffer (20 mM BTP, pH 7.9, 10% glycerol, 35 mM NaCl, 1 mM TCEP) and incubated for 6 h. The flow-through was collected and used to purify rhodopsin or PRPH2/ROM1. After washing the column with 15 mL of SMA Wash Buffer, two successive 15 mL washes with High Salt SMA Wash Buffer (20 mM BTP, pH 7.9, 10% glycerol, 500 mM NaCl, 1 mM TCEP) were passed over the column, followed by a 15 mL wash with SMA Wash Buffer. Elution Buffer was made by adding 40 mg/mL of CL2 peptide (NETYDLPLHPRTAGASRQAKEVDKGC) to 1 mL of Wash Buffer. After the elution step, the column was washed with 1 mL of SMA Wash Buffer, and then all proteins remaining on the resin were eluted with 1 column volume of 10% SDS. Each lane of the corresponding SDS-PAGE gel represents 10 μL of sample at the concentration of the sample, not adjusted to constant protein concentration across lanes.

Immunoaffinity Elution and Elution Wash fractions of ABCA4 were pooled and concentrated to 0.5 mL and then centrifuged at 20,000*g* for 10 min. The soluble fraction was then injected onto a Superdex 200 Increase 10/300 GL (GE Healthcare Bio-Sciences) size exclusion chromatography (SEC) column to remove rhodopsin. SMA SEC Buffer (20 mM BTP, pH 7.9, 10 mM NaCl, 1 mM TCEP) was used as the mobile phase, and fractions containing ABCA4 were pooled for use in other experiments.

### Establishing Nb for PRPH2/ROM1 isolation

Washed ROS membranes from 50 frozen bovine retinas were thawed on ice and re-suspended in a detergent-based solubilization buffer (20 mM BTP, pH 7.9, 300 mM NaCl, 2.5 mM DTT, 25 mM DDM) and incubated at 4 °C for 1 h with end over end mixing. To prevent reactions between free cysteine residues, the crude protein lysate was treated with 5.0 mM iodoacetamide for 30 min at room temperature. The solution was then quenched with an additional 5 mM DTT and immediately centrifuged at 150,000 × *g* for 1 h at 4 °C to clear insoluble material and aggregated proteins, the sample then was incubated for 1 h at 4°C with end over end mixing with nanobodies Nb20, Nb19, Nb28, Nb32, Nb13 to a final ratio of PRPH2/ROM1:Nb of 1:2. 1.0 mL of pre-equilibrated cOmplete Ni^2+^-resin (Sigma-Aldrich) was added to the solution and incubated for 1 h at 4 °C with end-over-end mixing. The resultant suspension was transferred to a 5.0 mL gravity column. The resin was washed with 10 column volumes of 20 mM BTP, pH 7.9, 300 mM NaCl, 0.35 mM DDM, 1.0 mM imidazole. Each PRPH2/ROM1/Nb complex was eluted with four column volumes elution buffer, comprised of the same wash buffer but with a final imidazole concentration of 300 mM. Aliquots of all samples along the stages of purification were saved for analysis. The resulting elution was then concentrated, and buffer exchanged to 20 mM BTP pH 7.9, 300 mM NaCl, 0.35 mM DDM using a PD-10 column (GE Healthcare Bio-Sciences). The sample was concentrated to 1.0 mg/mL, frozen in liquid nitrogen, and stored at −80 °C for future use.

### Purification of native, bovine PRPH2/ROM1 in SMA

ROS isolated from 50 bovine retinas were thawed on ice and re-suspended in Extraction Buffer with 2.5% SMA (v/v) (XIRAN SL30010 P20) (Polyscope Polymers B.V.) and incubated at 4 °C for 1 h with end-over-end mixing. To prevent reactions between free cysteine residues, the crude protein lysate was treated with 5.0 mM iodoacetamide for 30 min at room temperature. The solution was then quenched with an additional 5 mM DTT and immediately centrifuged at 150,000 × *g* for 1 h at 4 °C to clear insoluble material and aggregated proteins. The sample then was incubated for 1 h at 4 °C with end-over-end mixing with PRPH2/ROM1-specific nanobody Nb19 to a final ratio of PRPH2/ROM1:Nb19 at 1:2 (Nb19 includes His6 tag). 1.0 mL of pre-equilibrated cOmplete Ni^2+^-resin (Sigma-Aldrich) was added to the solution and incubated for 1 h at 4 °C with end-over-end mixing. The resultant suspension was transferred to a 5.0 mL gravity column. The resin was washed with 10 column volumes of 20 mM BTP, pH 7.9, 300 mM NaCl, and 1.0 mM imidazole. The Prph2/ROM1/Nb19 complex was eluted with four column volumes of elution buffer, comprised of the same wash buffer but with a final imidazole concentration of 300 mM. Aliquots of all samples along the stages of the purification were saved for analysis. The resulting elution was then concentrated, and buffer exchanged to 20 mM BTP pH 7.9, 300 mM NaCl using a PD-10 column (GE Healthcare). The sample was concentrated to 1.0 mg/mL, frozen in liquid nitrogen, and stored at −80 °C for future use.

### Purification of native, bovine rhodopsin in SMA

ROS isolated from 50 bovine retinas were extracted in 16 mL of Extraction Buffer with 2.5% SMA (v/v) (XIRAN SL30010 P20) (Polyscope Polymers B.V.) for 1 h at 4 °C in the dark, followed by centrifugation at 100,000*g* for 1 h at 4 °C. 1 mL of ∼8.0 mg/mL fresh immunoaffinity resin was prepared by conjugating purified, anti-rhodopsin antibody (1D4) to CNBr-activated Sepharose 4B beads (GE Healthcare Bio-Sciences) according to manufacturer’s instructions (Molday and Molday, 2014). The extracted fraction of ROS was then mixed with the immunoaffinity resin, brought to 168 mM NaCl through dilution with SMA Wash Buffer (20 mM BTP, pH 7.9, 10% glycerol, 35 mM NaCl, 1 mM TCEP) and incubated for 6 h. The flow-through was collected and used to purify PRPH2/ROM1. After washing the column with 15 mL of SMA Wash Buffer, two successive 15 mL washes with High Salt SMA Wash Buffer (20 mM BTP, pH 7.9, 10% glycerol, 500 mM NaCl, 1 mM TCEP) were passed over the column, followed by a 15 mL wash with SMA Wash Buffer. Elution Buffer was made by adding 40 mg/mL of 1D4 peptide (TETSQVAPA) to 1 mL of Wash Buffer (Molday and Molday, 2014). After the elution step, the column was washed with 1 mL of SMA Wash Buffer, and then all proteins remaining on the resin were eluted with 1 column volume of 10% SDS. Each lane of the corresponding SDS-PAGE gel represents 10 μL of sample at the concentration of the sample, not adjusted to constant protein concentration across lanes.

### Transmission electron microscopy

Four μL of the peak SEC fractions containing ABCA4 were adsorbed for 1 min to carbon coated, glow-discharged grids (15 mA for 15 sec) (Electron Microscopy Sciences, Hatfield, PA). The grids were washed with two 20 μL drops of ultrapure water and then stained with two 20 μL drops of 1% (w/v) uranyl acetate (Electron Microscopy Sciences); the first for 10 sec and the second for 1 min. Data were collected with a JEOL JEM-2200fs microscope (JEOL, Japan), operated at 200 kV and equipped with a Tietz TVIPS CCD Camera at 60,000x magnification. The pixel size was 2.131 Å.

### Single particle reconstruction

*De novo* particle reconstruction of SMALP-imbedded ABCA4 was done using the program cisTEM. following a published workflow (Grant et al., 2018). cisTEM auto-picked 71,088 particles that were then sorted by 2D classification into good classes containing 14,652 particles. The particles contained in these classes were then used for cisTEM’s *ab initio* 3D structure generation, which was then refined using cisTEM’s Auto Refine. Further structural analysis of ABCA4 was done in UCSF Chimera (Pettersen et al., 2004).

### Trp fluorescence quenching assay

All measurements were performed on a PerkinElmer Life Sciences LS55 model fluorometer (PerkinElmer, Waltham, MA). Binding of ATP to purified ABCA4 in SMALPs was evaluated by monitoring the quenching of protein fluorescence at increasing concentrations of ATP (0-1.5 mM). With the excitation wavelength set at 290 nm, emission spectra were recorded at 330 nm over 1 min with 2 sec intervals with bandwidths for excitation and emission fixed at 10 nm. Titrations were carried out at 20 °C in 20mM BTP buffer, pH 7.9, containing 35 mM NaCl and 1 mM TCEP. ATP stock solution was diluted in ultrapure water. All binding data were corrected for background and self-absorption of excitation and emission light using a Varian Cary 50 Bio UV-Visible Spectrophotometer (Palo Alto, CA).

### Rhodopsin absorption assay

All measurements were performed on a Varian Cary 50 Bio UV-Visible Spectrophotometer. Rhodopsin purified in the dark in SMALPs was measured by absorption from 250-600 nm. The sample was then incubated with hydroxylamine to a final concentration of 8 mM and allowed to bleach completely in light for 7 min, after which the absorption spectrum was taken. The sample was regenerated with 9-cis retinal added to a final concentration of 70 μM and allowed to regenerate over 20 min, overnight, and for two days, with the spectrum taken at each time point.

### Lipid extraction and untargeted lipidomics

Lipids were extracted using a modified version of the Bligh-Dyer method (Bligh and Dyer, 1959). Briefly, samples were shaken in a glass vial (VWR) with 1 mL PBS, 1 mL methanol and 2 mL chloroform containing internal standards (^13^C_16_ palmitic acid, ^2^H_7_ cholesterol) for 30 sec. The resulting mixture was vortexed for 15 sec and centrifuged at 2400*g* for 6 min to achieve phase separation. The organic (bottom) layer was retrieved using a Pasteur pipette, dried under a gentle stream of nitrogen, and reconstituted in 2:1 chloroform:methanol for LC/MS analysis.

Lipidomic analysis was performed on a Vanquish HPLC online with a Q-Exactive quadrupole-orbitrap mass spectrometer equipped with an electrospray ion source (Thermo). Data was acquired in positive and negative ionization modes. Solvent A consisted of 95:5 water:methanol, Solvent B was 70:25:5 isopropanol:methanol:water. For positive mode, solvents A and B contained 5 mM ammonium formate with 0.1% formic acid; for negative mode, solvents contained 0.028% ammonium hydroxide. An XBridge (Waters) C8 column (5 μm, 4.6 mm × 50 mm) was used. The gradient was held at 0% B between 0 and 5 min, raised to 20% B at 5.1 min, increased linearly from 20% to 100% B between 5.1 and 55 min, held at 100% B between 55 min and 63 min, returned to 0% B at 63.1 min, and held at 0% B until 70 min. Flow rate was 0.1 mL/min from 0 to 5 min, 0.3 mL/min between 5.1 min and 55 min, and 0.4 mL/min between 55 min and 70 min. Spray voltage was 3.5 kV and 2.5 kV for positive and negative ionization modes, respectively; S-lens RF level was 65. Sheath, auxiliary, and sweep gases were 50, 10 and 1, respectively.

Capillary temperature was 325 °C and auxiliary gas heater temperature was 200 °C. Data were collected in full MS/dd-MS2 (top 10). Full MS was acquired from 150–1500 m/z with resolution of 70,000, AGC target of 1×10^6^ and a maximum injection time of 100 ms. MS2 was acquired with resolution of 17,500, a fixed first mass of 50 m/z, AGC target of 1×10^5^ and a maximum injection time of 200 ms. Stepped normalized collision energies were 20, 30 and 40%.

### Lipid extraction and FA lipidomic analysis

For lipid hydrolysis, extracted lipids were resuspended in 200 μL of ethanol, incubated with 0.1 M KOH at room temperature for 24 h for saponification. The reaction was stopped by addition of 0.2 M HCl. Lipids were extracted as described above with ^2^H_31_ palmitic acid as internal standard.

FA lipidomic analysis was performed on a Dionex Ultimate 3000 LC system (Thermo) coupled to a TSQ Quantiva mass spectrometer (Thermo). Solvent A consisted of 95:5 water:methanol, Solvent B was 70:25:5 isopropanol:methanol:water. For negative mode, solvents contained 0.028% ammonium hydroxide. An XBridge C8 column (Waters, Milford, MA) (5 μm, 4.6 mm × 50 mm) was used. The gradient was as described under “**Lipid extraction and untargeted lipidomics**” (above). MS analyses were performed using electrospray ionization in negative ion mode, with spay voltages of −2.5 kV, ion transfer tube temperature of 325 °C, and vaporizer temperature of 200 °C. Sheath, auxiliary, and sweep gases were 40, 10 and 1, respectively. Pseudo-MRM was performed for all fatty acids.

### Lipid data analysis

Lipid identification was performed with LipidSearch (Thermo). Mass accuracy, chromatography and peak integration of all LipidSearch-identified lipids and targeted lipids were verified with Skyline (MacLean et al., 2010). Peak areas were used in data reporting, and data were normalized using internal standards. Quantification of the FFAs was performed by measuring the area under the peak, and the “raw” value is reported as relative molar percentage of total area under the curve for each sample. In cases of two peaks for a single species (*e.g.*, the result of omega-3 vs omega-6 differences in FA), we added the peak areas together and reported the species without omega-3/6 differentiation. Each lipid class was then normalized separately such that the sum of all species of a class equaled 100%. These relative molar percentages were used for all graphs and analysis. In cases of less than 3 samples for a particular species, the species was excluded from all ANOVA analysis. All lipid species found across all samples were used for PCA (199 total species, 14 classes). PCA scores, loadings, and variances were calculated using Graphpad Prism software (Graphpad, San Diego, CA).

## SUPPLEMENTAL MATERIAL

**Fig. S1** shows SMA extraction of ROS, characterization of anti-ABCA4 mAb CL2 with murine and bovine samples, and nsTEM analysis showing ABCA4 purified with CL2 in SMA shows increased TMD density.

**Fig. S2-5** show the full list of lipid species detected, with each species amount graphed as the percent of the total for each particular class (PC, PE, etc.).

## ACKNOWEDGEMENTS

We would like to thank David Peck, Tim Dinh, and Huajun Yan for isolation of the bovine retinas. We would also like to thank Brian Kevany for help with the initial nanobody expression and screening The CL2 antibody was produced by Denice Major, Visual Science Research Core of Case Western Reserve University, supported by P30 EY11373. This research was supported in part by grants to K.P. from the National Institutes of Health (NIH) (EY009339, EY027283, EY030873, and EY019312) and to P.D.K. from the U.S. Department of Veterans Affairs (I01BX004939). C.L.S. was supported by NEI-funded predoctoral fellowships T32EY007157-17 and T32EY007157-16A1. E.H.C. was supported by predoctoral fellowships T32GM007250 and T32GM008803. S.S. was supported by predoctoral fellowships F30EY029136-01A1, T32EY024236, and T32GM007250. The authors also acknowledge support from an RPB unrestricted grant to the Department of Ophthalmology, University of California, Irvine. This work was also supported by the Mass Spectrometry Core of the Salk Institute with funding from NIH-NCI CCSG: P30 014195 and the Helmsley Center for Genomic Medicine. The MS data described here were gathered on a ThermoFisher Q Exactive Hybrid Quadrupole Orbitrap mass spectrometer funded by NIH grant (1S10OD021815-01). Molecular graphics and analyses were performed with UCSF Chimera, developed by the Resource for Biocomputing, Visualization, and Informatics at the University of California, San Francisco, with support from NIH P41-GM103311. C.L.S., H.J., and K.P. have filed for a patent on the CL2 monoclonal antibody; they declare no additional conflict of interest. All other authors report no conflicts of interest.

## AUTHOR CONTRIBUTIONS

Christopher L. Sander helped design the antigen for the CL2 mAB; designed, performed, and/or analyzed the results of all experiments; and wrote and revised the manuscript. Avery E. Sears developed the Nb19 nanobody and the purification of PRPH2/ROM1 and helped write and revise the manuscript. Antonino M. Pinto and Alan Saghatelian helped design and carried out the lipidomic data collection and performed initial data analysis. Elliot H. Choi performed IHC of murine samples and revised the manuscript. Shirin Kahremany helped design and perform rhodopsin regeneration experiments and revised the manuscript. Susie Suh performed immunoblots of murine samples and revised the manuscript. Hui Jin helped design the antigen for the CL2 mAB and revised the manuscript. Els Pardon and Jan Steyaert developed and provided the original Nb families for screening against PRPH2/ROM1 and helped edit the manuscript. Zhiqian Dong provided mouse retina cryosections. Dorota Skowronska-Krawczyk helped design the lipidomic experiments and revised the manuscript. Philip D. Kiser helped design all experiments and revised the manuscript. Krzysztof Palczewski helped design the antigen for the CL2 mAb, helped design all experiments, and revised the manuscript.

## ABBREVIATIONS

The abbreviations used are

ABC: ATP-binding cassette
AcCa: acylcarnitine
AMP: adenosine monophosphate
ATR: all-trans retinal
BTP: bis-tris propane
CDR: complimentary determining regions
Cer: ceramides
ChE: cholesterol ester
CHS: cholesterol hemisuccinate
CMC: critical micelle concentration
CNBr: cyanogen bromide
cryoTEM: cryogenic transmission electron microscopy
DDM: n-dodecyl â-D-maltoside
DG: diacylglycerol
DHA: docosahexaenoic acid
DIBMA: diisobutylene maleic acid
DIBMALP: diisobutylene maleic acid lipid particles
ECD: extracytosolic domain
EW: elution wash
FA: fatty acid
FFA: free fatty acid
FT: flow-through
FWR: framework regions
GPCR: G protein-coupled receptor
H1/2/3: hypervariable regions or loops 1/2/3
IHC: immunohistochemistry
kDa: kilodalton
KLH: keyhole limpet hemocyanin
KO: knock-out
L: load
LC-MS: liquid chromatography-mass spectrometry
LC-PUFA: long chain-polyunsaturated fatty acid
LMNG: laurel maltose neopentyl glycol
LPA: lyso-phosphatidic acid
LPC: lyso-phosphatidylcholine
LPE: lyso-phosphatidylethanolamine
LUV: large unilamillar vesicles
lyso-PL: lyso-phospholipid
mAb: monoclonal antibody
MG: monoacylglycerol
MS: mass spectroscopy
MSP: membrane scaffold protein
Nb: nanobody
N-ret-PE: N-retinylidene-phosphatidylethanolamine
nsTEM: negative stain transmission electron microscopy
PA: phosphatidic acid
PAGE: polyacrylamide gel electrophoresis
PBS: phosphate-buffered saline
PBST: phosphate-buffered saline with Tween-20
PC: phosphatidylcholine
PE: phosphatidylethanolamine
PRPH2: peripherin2
PI: phosphatidylinositol
PS: phosphatidylserine
PVDF: polyvinylidene difluoride
Res: resin
ROM1: rod outer segment membrane protein 1
ROS: rod outer segment
RPE65: retinal pigment epithelium-specific 65 kDa protein
RT: room temperature
SEC: size exclusion chromatography
SMA: styrene maleic acid
SMALP: styrene maleic acid lipid particle
SDS: sodium dodecyl sulfate
TCEP: tris(2-carboxyethyl)phosphine)
TG: triacylglycerol
TMD: transmembrane domain
VLC-PUFA: very long chain-polyunsaturated fatty acid
W1-4: wash 1-4
WT: wild-type

## Supplemental Figures

**Supplemental Figure 1.**
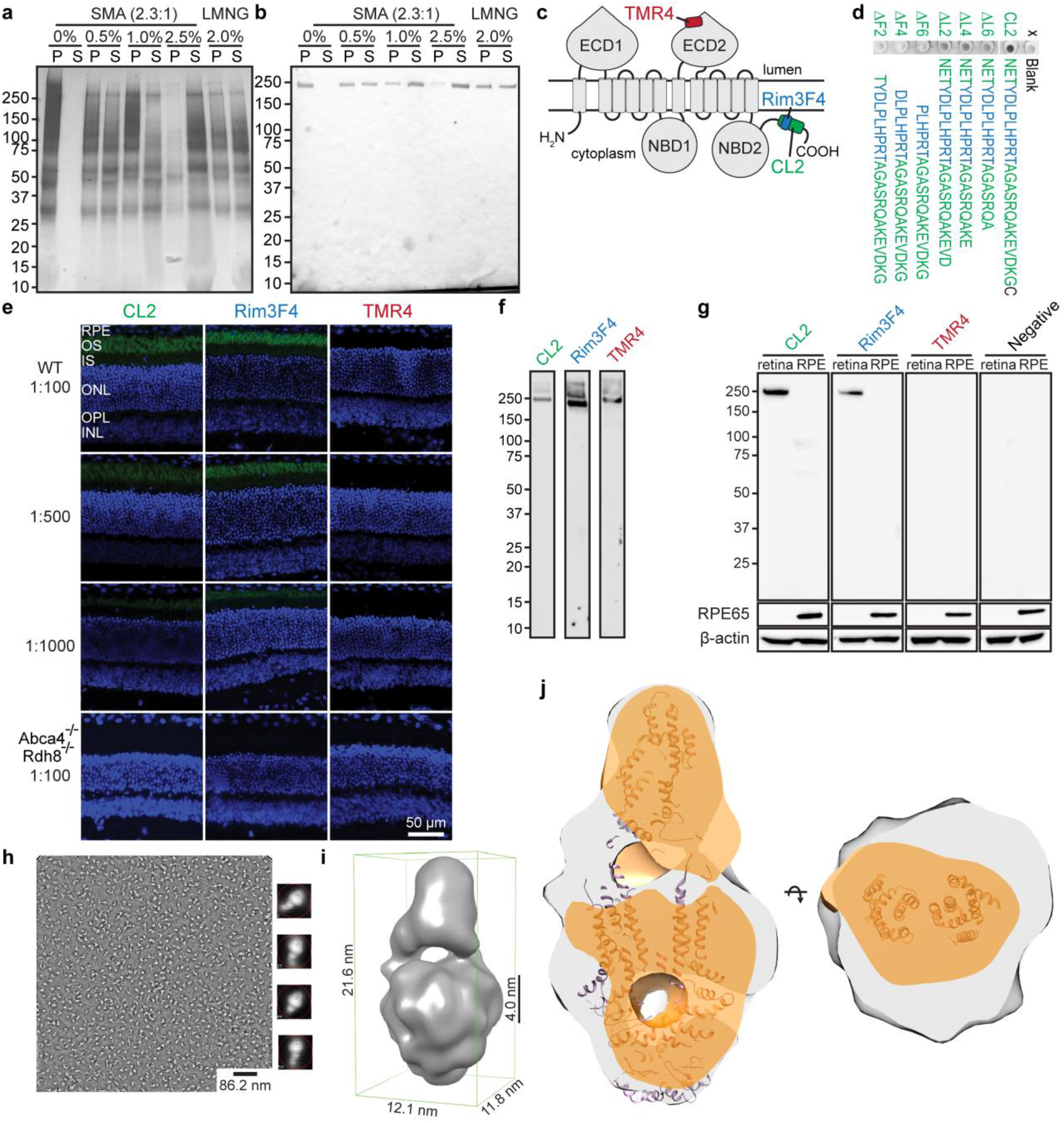
SMA extracts ROS yielding more ABCA4, mAb CL2 recognizes ABCA4, and ABCA4 purified with CL2 shows increased transmembrane density, suggesting SMALP has formed. (**a**) Extraction of ROS proteins by various concentrations of SMA, or by the low-CMC detergent LMNG. Residual ROS pellets after initial detergent extraction were solubilized with 10% SDS. P, pellet; S, soluble. (**b**) Immunoblotting demonstrates a graded extraction of ABCA4 with increasing amounts of SMA. (**c**) Topographical map of ABCA4 highlighting the epitopes of three monoclonal antibodies, TMR4, Rim3F4, and CL2. (**d**) Dot blots of polypeptides comprised of the amino acid chains shown to the right were used to confirm the novel epitope of CL2 on the C-terminus of ABCA4. Truncations of the beginning of the sequence decreased the binding of CL2. The Rim3F4 epitope is depicted in blue. (**e**) Immunohistochemistry of retinal cryosections from 2-month-old WT and *Abca4*^-/-^*Rdh8*^-/-^ KO mice, using CL2, Rim3F4 and TMR4 antibodies against ABCA4 (green) at three different dilutions. As expected, no fluorescence signal occurred with the KO mouse cryosections. With cryosections from WT mice, primary incubations with CL2 and Rim3F4 antibodies showed specific immunoreactivity with photoreceptor outer segments at all three dilutions, whereas TMR4 did not generate a fluorescence signal. Scale bar: 50 µm. (**f**) Relative amount of ABCA4 present in solubilized bovine ROS as assessed by immunoblotting. Stock concentrations of 1 mg/mL were used for all antibodies, and the dilution aw 1:10,000 for each antibody tested. (**g**) Immunoblot of retinal and RPE lysates obtained from 2-month-old WT mouse using CL2, Rim3F4 and TMR4 antibodies. Probing with CL2 and Rim3F4 antibodies resulted in a specific band at 250 kDa in the retinal samples, which corresponds to the size of ABCA4; whereas no positive signal was detected with TMR4. RPE65 (65 kDa) served as the control for tissue sample purity, and β-actin (42 kDa) served as the loading control. (**h**) Negative stain micrograph of a representative SMA-CL2 preparation with 2D classes to the right; 60,000x magnification. SMALP-extracted ABCA4 shows an increase in transmembrane domain density, indicative of a native lipid belt. Scale bar: 86.2 nm. (**i**) 3D reconstruction of ABCA4 at ∼ 18 Å resolution showing a putative bilayer thickness in the region of the SMALP. (**j**) SMALP-imbedded ABCA4 (gray) shows considerably more density within the predicted TMD region compared to: (1) a prior ABCA4 negative-stained structure (EMDB-5497 (orange), solubilized in n-dodecyl β-D-maltoside (DDM) and then switched into amphipol); and (2) the ABCA4 homolog, ABCA1 (EMDB-6724 (purple ribbon), solubilized in DDM and cholesterol hemisuccinate (CHS) and then switched into digitonin). We interpret these differences to be explained by the SMALP nanodisc containing native lipids surrounding the TMD of ABCA4.

**Supplemental Figure 2.**
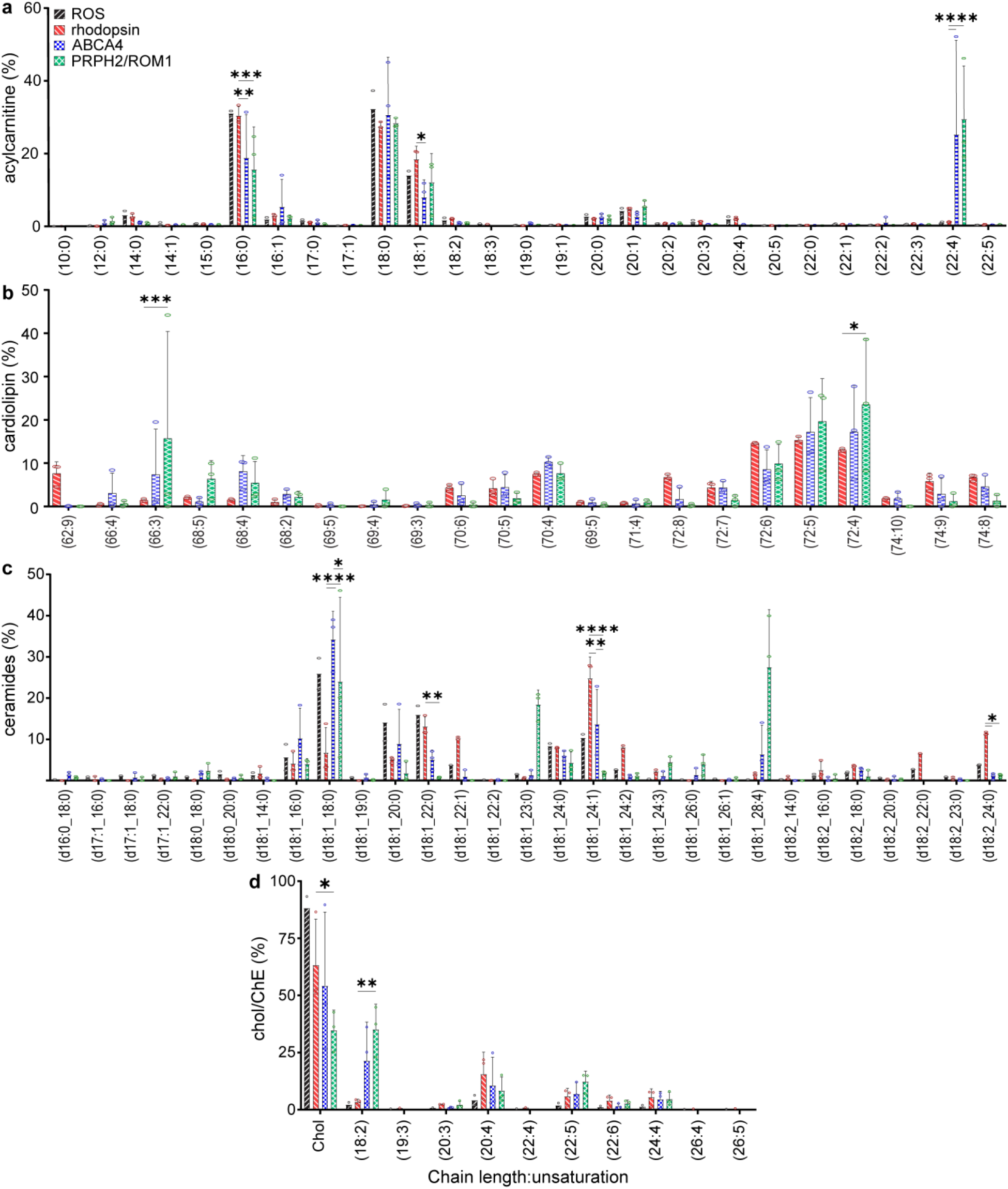
Complete set of detected species for AcCa, cardiolipin, Cer, and cholesterol/ChE. **(a-d)** Every detected species of lipid that copurified with each sample is shown as a percentage of each respective class (class noted on y-axis). Cardiolipin chain lengths and unsaturation levels summed together. Total ROS: black forward stripe; rhodopsin: red backward stripe; ABCA4: blue checker; PRPH2/ROM1: green diamond. Number of measurements for each sample of each species varies and is noted by the individual data points for each bar (open circles). Percent composition was calculated for each sample by dividing the area under the curve for each species in a class by the total area under the curve for that class, measured *via* LC-MS after correction for variations in internal standard area, sample mass, and sample injection volume. Statistics were determined using two-way ANOVA with Tukey’s multiple comparisons post-hoc test between samples that had at least 3 detected replicates. Statistical significance values are indicated as follows: *, *P*<0.05; **, *P*<0.01; ***, *P*<0.001; ****, *P*<0.0001.

**Supplemental Figure 3.**
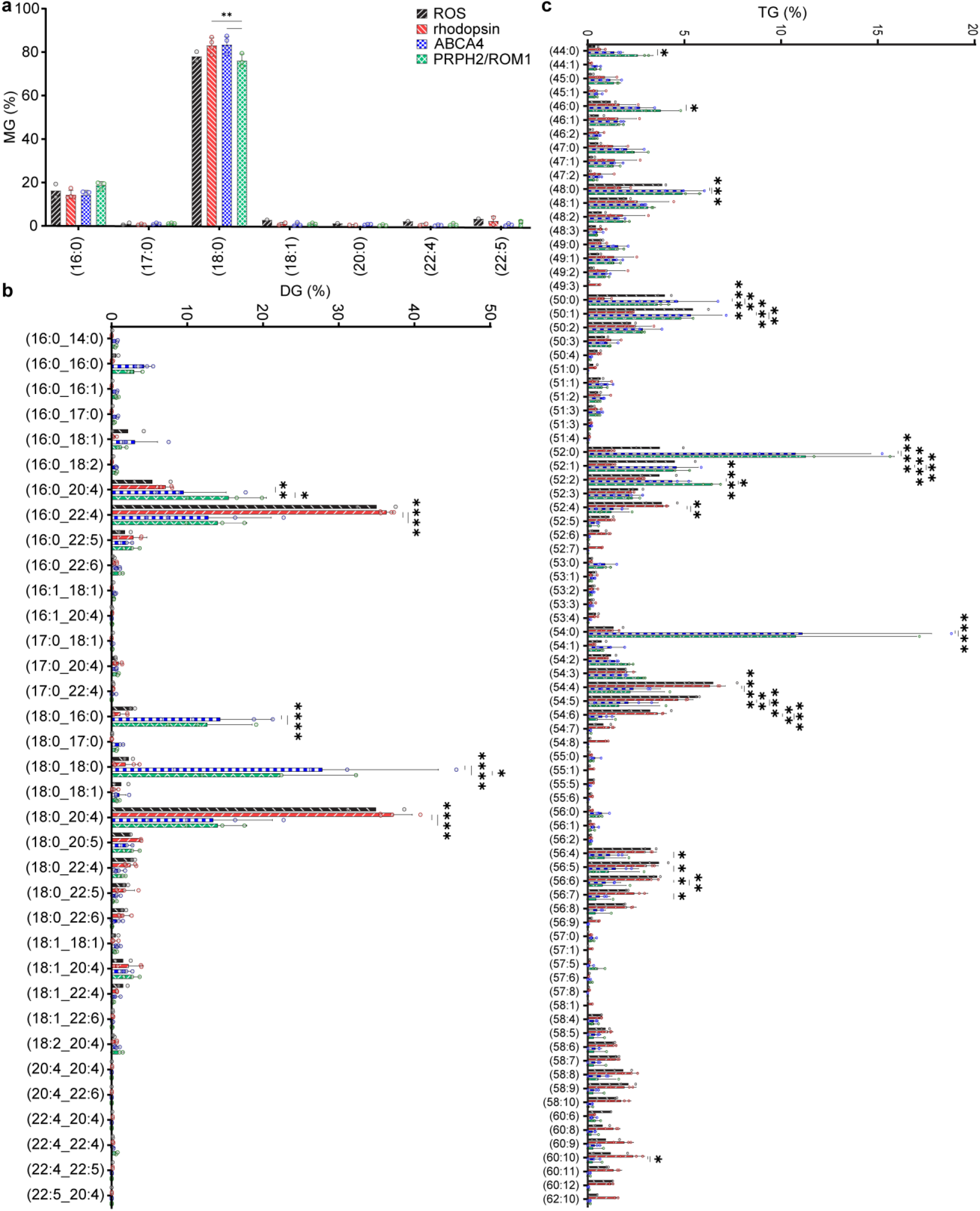
Complete set of detected species for mono-, di-, and triacylglicerides. **(a-c)** Every detected species of lipid that copurified with each sample is shown as a percentage of each respective class (class noted on y-axis). Triacylgliceride chain lengths and unsaturation levels summed together. Total ROS: black forward stripe; rhodopsin: red backward stripe; ABCA4: blue checker; PRPH2/ROM1: green diamond. Number of measurements for each sample at each species varies, and is noted by the individual data points for each bar. Percent composition was calculated for each sample by dividing the area under the curve for each species in a class by the total area under the curve for that class, measured *via* LC-MS after correction for variations in internal standard area, sample mass, and sample injection volume. Statistics are determined using two-way ANOVA with Tukey’s multiple comparisons post-hoc test between samples that had at least 3 detected replicates. Statistical significance values are indicated as follows: *, *P*<0.05; **, *P*<0.01; ***, *P*<0.001; ****, *P*<0.0001.

**Supplemental Figure 4.**
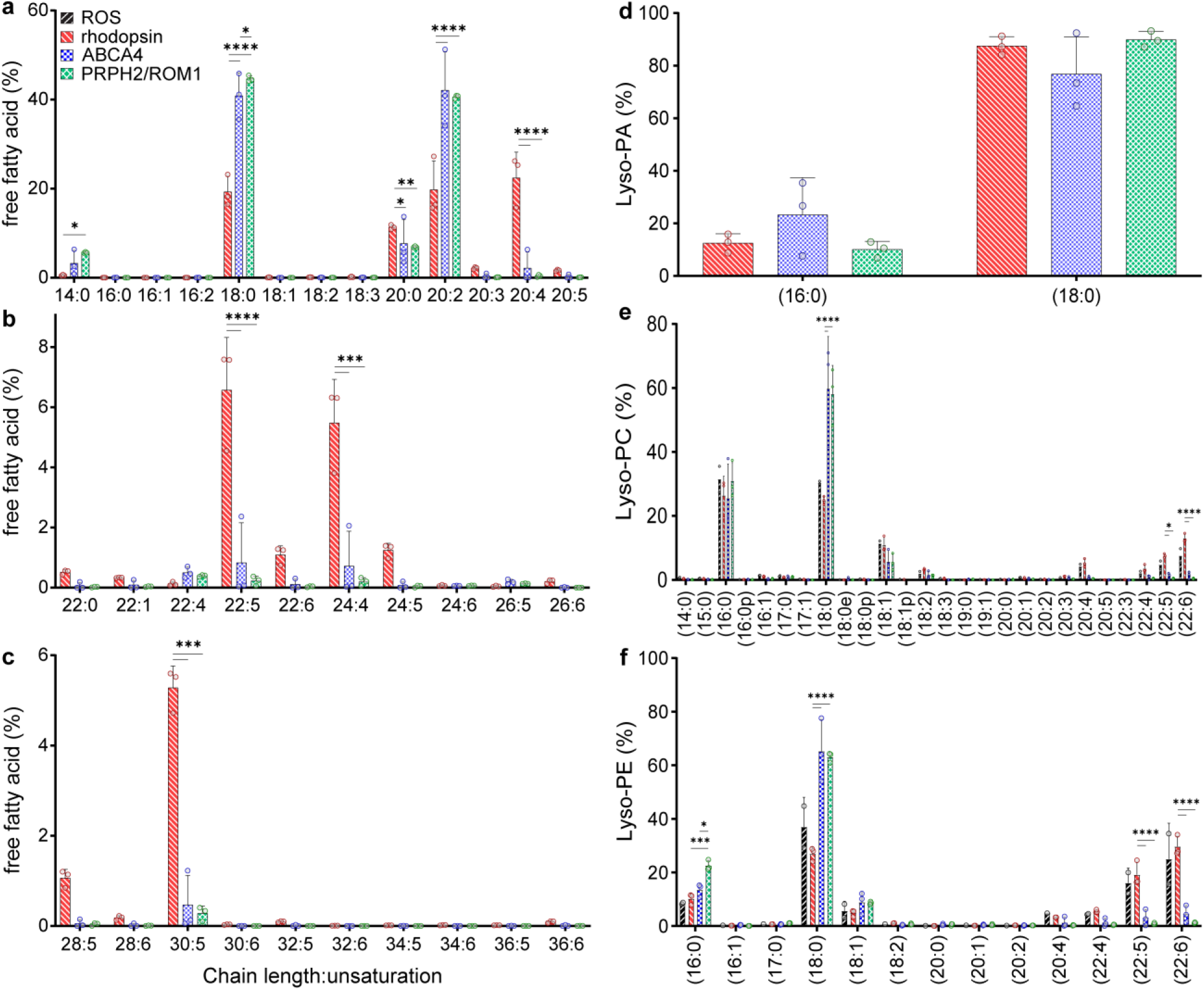
Every detected species of FFA and Lyso-PL, relative to total species of each class. **(a-f)** Every detected species of FFA and lyso-PL that copurified with each sample is shown as a percentage of each respective class (class noted on y-axis). Total ROS: black forward stripe; rhodopsin: red backward stripe; ABCA4: blue checker; PRPH2/ROM1: green diamond. Number of measurements for each sample at each species varies, and is noted by the individual data points for each bar (open circles). Percent composition was calculated for each sample by dividing the area under the curve for each species in a class by the total area under the curve for that class, measured *via* LC-MS after correction for variations in internal standard area, sample mass, and sample injection volume. Statistics were determined using two-way ANOVA with Tukey’s multiple comparisons post-hoc test. Statistical significance values are indicated as follows: *, *P*<0.05; **, *P*<0.01; ***, *P*<0.001; ****, *P*<0.0001.

**Supplemental Figure 5.**
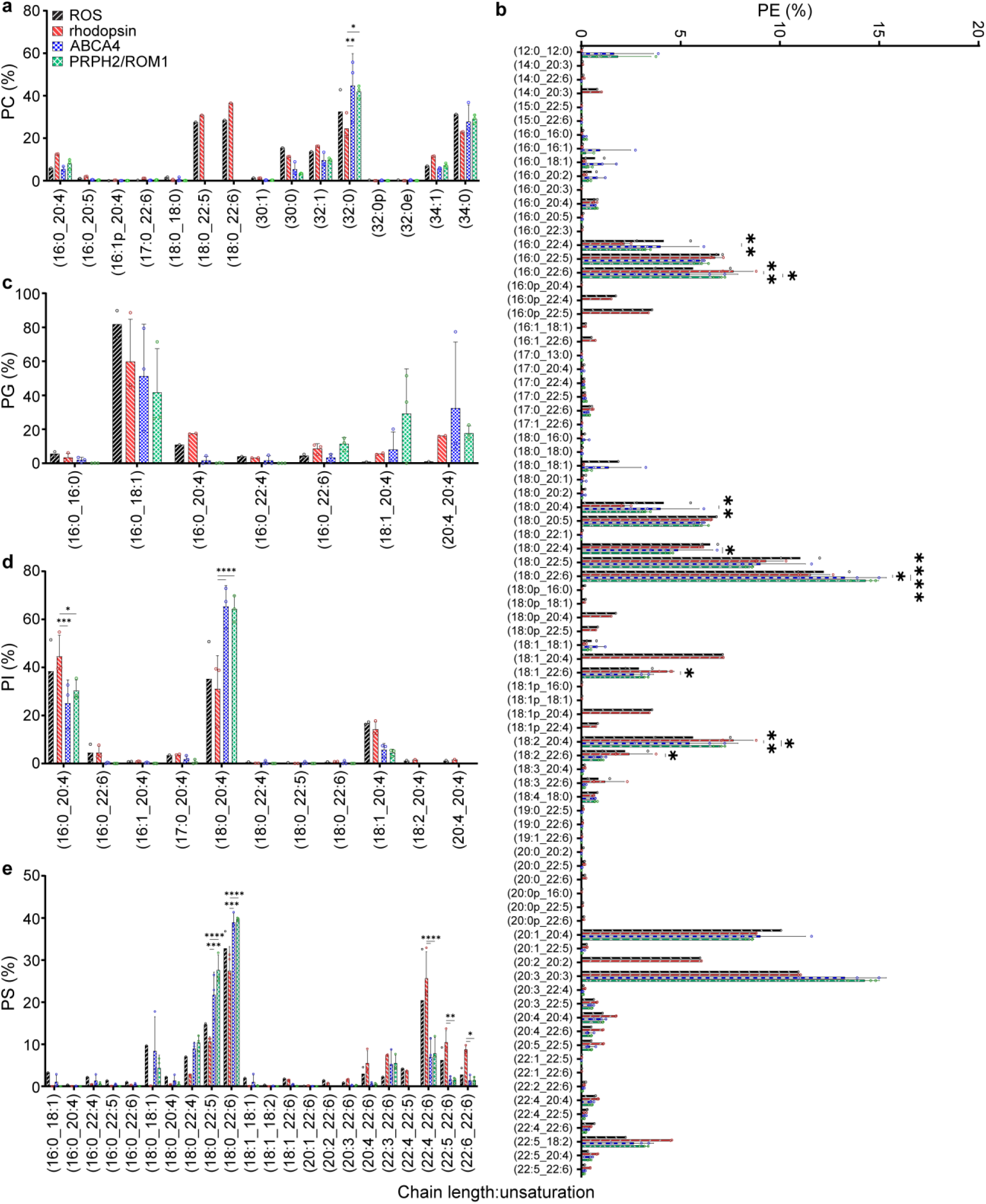
Every detected species of PL, relative to total species of each class. **(a-e)** Every detected species of PL that copurified with each sample is shown as a percentage of each respective class (class noted on y-axis). Total ROS: black forward stripe; rhodopsin: red backward stripe; ABCA4: blue checker; PRPH2/ROM1: green diamond. Number of measurements for each sample at each species varies, and is noted by the individual data points for each bar (open circles). Percent composition was calculated for each sample by dividing the area under the curve for each species in a class by the total area under the curve for that class, measured *via* LC-MS after correction for variations in internal standard area, sample mass, and sample injection volume. Statistics were determined using two-way ANOVA with Tukey’s multiple comparisons post-hoc test. Statistical significance values are indicated as follows: *, *P*<0.05; **, *P*<0.01; ***, *P*<0.001; ****, *P*<0.0001.

